# Psoriasis-related neuroinflammation disrupts thalamostriatal signalling driving anhedonia in both humans and mice

**DOI:** 10.1101/2024.06.26.600791

**Authors:** Deepika Sharma, Lilya Andrianova, Rhona McGonigal, Kirstyn Gardner-Stephen, Hassan al Fadhel, Flavia Sunzini, Kristian Stefanov, Dominique L Chaput, Jennifer A Barrie, Richard Hohne, Megan Saathoff, Yaprak Karabalci, Alfredo Montes Gomez, Julie-Myrtille Bourgognon, Ana González-Rueda, Peter Thornton, Neil Basu, John J Cole, Michael T Craig, Jonathan T Cavanagh

## Abstract

Inflammation is implicated in 25% of depression cases, yet limited access to human brain for mechanistic studies and scarce translational models have hindered the identification of neural circuits linking systemic inflammation to depressive symptoms such as reduced motivation and anhedonia. Leveraging both clinical and pre-clinical approaches, we combined neuroimaging in individuals with psoriatic disease, a systemic inflammatory condition frequently associated with depression, with neurophysiological, behavioural and immunological studies in a psoriasis mouse model exhibiting neuroinflammation. We found that increased inflammation had a significant association with both depression and fatigue in individuals with psoriatic disease. Across species, we found that inflammatory signalling disrupts thalamostriatal circuitry, a key component of the motivational network. In humans, functional connectivity between thalamus and ventral striatum correlated with depressive and fatigue-related symptoms. In mice, psoriasis-like inflammation produced impaired thalamostriatal synaptic transmission accompanied by anhedonia and motivation-related behavioural deficits, together with glial activation and immune-cell infiltration. These cross-species findings identify the thalamostriatal circuit as a conserved neuroinflammatory hotspot involved in depression and highlight it as a potential therapeutic target.

## Introduction

Neuroinflammation is emerging as a key driver in the pathophysiology of depression and other neuropsychiatric disorders: up to 75% of patients with immune-mediated inflammatory diseases (IMIDs) such as psoriatic disease (PsD) experience comorbid depression^1,2^. Targeted anti-inflammatory biologic therapies can alleviate this comorbid depression in IMIDs^3^, and a recent meta-analysis linked increased peripheral inflammation to antidepressant resistance^4^. Meta-analyses of multiple GWAS studies have implicated immune-related genes in MDD^5^, and numerous studies have linked exogenous cytokine administration to disrupted neural circuits and networks key to motivation, threat detection and emotional processing^6,7^. As part of the brain’s reward network, the cortico-thalamostriatal circuitry has been consistently linked to motivational anhedonia, strongly associated with MDD^8^. On this basis, we hypothesised that thalamostriatal circuit perturbations would be evident in both humans and mice in the context of inflammation, providing a firm translational basis for mechanistic studies of the role of immune activation in MDD.

Human brain imaging reveals that IFN-treatment rapidly reduced global connectivity within the brain, and local changes in connectivity between striatum and cortex^7^. Subsequent work found that treatment with IFN- and anti-TNF (tumour necrosis factor-) had opposing effects on depressive symptoms and resulted in consistent changes in brain connectivity^9,10^. Similarly, chronic mild stress in rats drives anhedonic behaviour and is associated with increased brain levels of TNFα and IL-1β^11^. Inflammation in mice leads to infiltration of peripheral monocytes into the brain and drives changes in both behaviour and dendritic morphology via TNFα-dependent mechanisms^12^. Both clinical and animal data show that the cellular sources of cytokines in the brain include both resident and recruited immune cells, with resident cell cytokine release modulated by the NLPR3 inflammasome^3^. TNF plays an important role in homeostatic control of glutamatergic neurotransmission, with astrocyte-derived TNF required both for scaling of synaptic strength^13^ and astrocytic glutamate cycling^14^. But excessive TNF can be maladaptive and can perturb striatal glutamatergic neurotransmission^15^. Application of TNF to brain slices evokes region-specific changes in synaptic function, reducing the AMPA/NMDA ratio in striatum but increasing it in hippocampus^16^. TNF acts directly on TNFR1 receptors on neurons to alter AMPA and GABA receptor trafficking within minutes of TNF application in culture^17^. Many of these effects are likely modulated by inflammation altering the ability of astrocytes to regulate synaptic levels of glutamate, with this mechanism increasingly being posited as a causal factor in mood disorders^18^.

To address our central hypothesis, we measured functional connectivity in the thalamostriatal network using resting state MRI in people with PsD and used the Aldara preclinical model of psoriasis (repeated topical application of the TLR7/8 agonist imiquimod over 3 days) to probe mechanistic changes in thalamostriatal circuitry and brain – immune interactions. Perturbation of thalamostriatal circuitry occurred in both clinical and preclinical studies and was associated with depression and deficits in motivational/rewarded behaviours. In the preclinical model, we found that reactive glia and infiltrating myeloid and lymphoid immune cells all made contributions to the neuroinflammatory environment within the brain, with the glial response mediated by the NLRP3 inflammasome. Our cross-species approach demonstrates the translational utility of studying thalamostriatal circuitry to find causal mechanisms linking neuroinflammation to deficits in motivation seen in depression.

## Results

### Depression and fatigue are associated with an inflammatory signature in individuals with psoriatic disease

In a cohort of individuals with PsD (see supplementary table 1 for clinical characterisation of our cohort), before treatment with biologic anti-cytokine therapy, we analysed peripheral blood for a large number of cytokines and other proteins (see methods). We carried out an exploratory analysis to determine whether the degree of inflammation experienced by these individuals was related to their depression or fatigue scores. Due to the nature of immune signalling in cascades, it was expected a high degree of collinearity in the blood markers, so we deployed an elastic net^19^ penalised regression model to determine, in an unbiased manner, whether any cytokines had a significant influence on either depression or fatigue scores. We found that no individual cytokine influenced either depression or fatigue scores. We employed a dimensional reduction approach and used a principal component analysis (PCA) of all blood markers and ran these in a linear model against both depression and figure. Using the complete blood marker panel, we found no significant relationship between blood markers and either depression (F=1.18, p=0.317) or fatigue (F=0.28, p=0.7566) in our cohort.

For an additional exploratory analysis, we carried out multiple correlations between each blood marker and both depression and fatigue scores (supplementary figure 1), excluding any nonsignificant correlations after correcting for multiple comparisons using the FDR. This exploratory analysis revealed that a small number of markers correlated with depression (positive correlation: BAFF, erythropoietin, CXCL1 and leptin; negative correlation: eotaxin and PYY) and fatigue (positive correlation: erythropoietin, IL1Ra, IL2Ra, CCL3 and leptin; negative correlation: eotaxin and IL17D). To determine whether these markers could explain any of the variability in depression and fatigue, we generated separate PCAs for depression and fatigue using just those biomarkers (loadings of the PCA are shown in figure 1A) and modelled the relationship between the first component of the PCA and these two measures in linear models. We found a significant relationship between the first principal component depression score (F=9.84; p=0.0031; figure 1B), with a moderate effect size (R^2^(adjusted) = 0.17). Similarly, we observed a significant relationship between the first principal component and fatigue score (F=9.1, p=0.0043; figure 1C) with a moderate effect size (R^2^(adjusted) =0.16). To control for the influence of lifestyle factors, we ran BMI and smoking status in linear models separately for depression and fatigue. We found that BMI, but not smoking status was a significant factor influencing variability in both depression and fatigue although the overall regression models fell short of formal statistical significance (Depression model: F=2.56, p=0.068, R^2^(adjusted) = 0.096; Fatigue model: F=2.59, p=0.066, R^2^(adjusted) = 0.098). Introducing BMI into the reduced biomarker model for depression led to an improved fit in the regression, with no significant effect of BMI alone in the model, but with a significant interaction between BMI and the first PCA of the biomarkers (overall model: F=6.2, p=0.0014, R^2^(adjusted) = 0.26). Including BMI in the fatigue model produced a small increase in R^2^(adjusted), to 0.19, but with no clear effect of either BMI, the PCA or their interaction apparent in the model.

**Figure 1:**
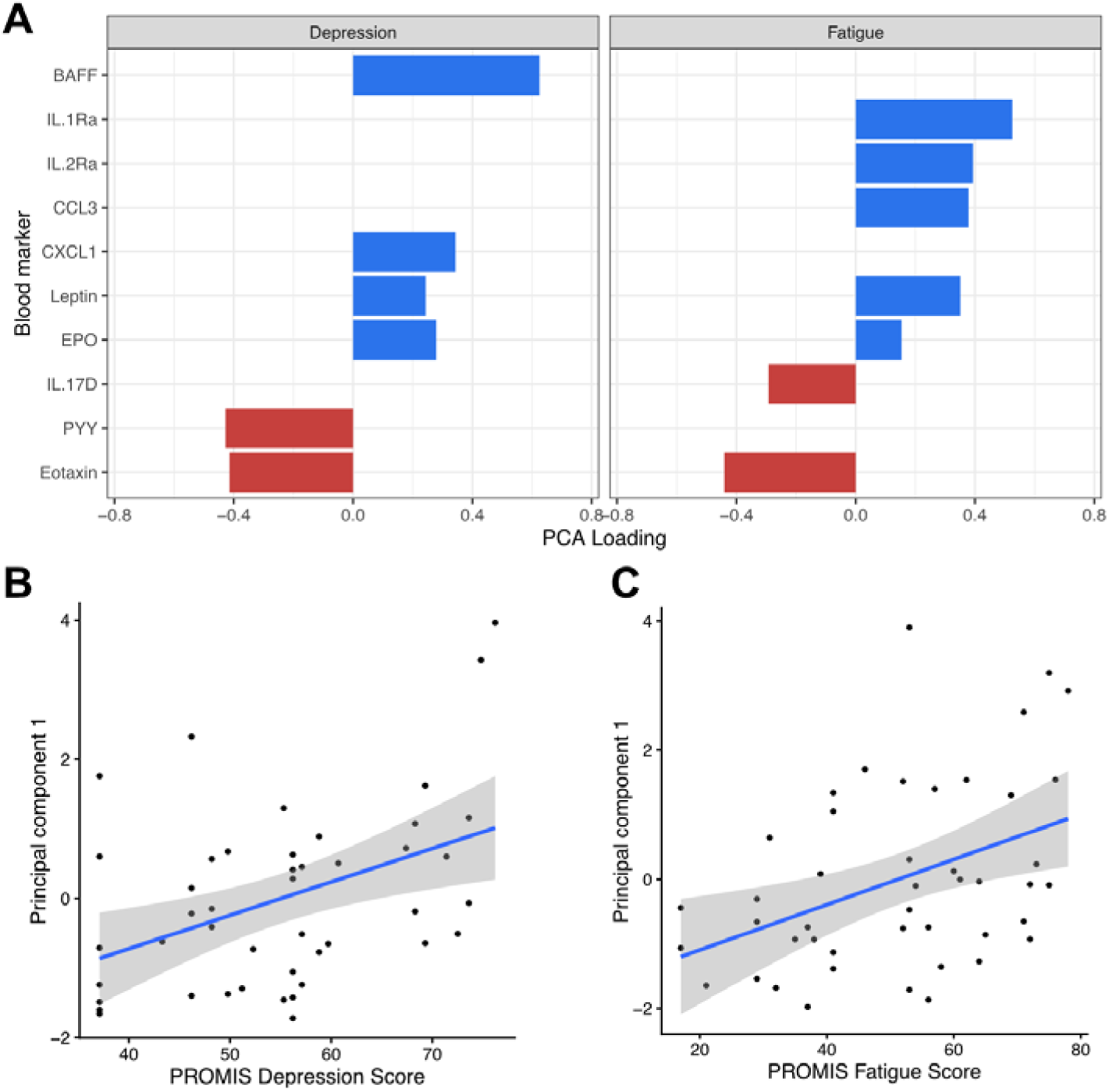
Relationship between blood markers, depression and fatigue. **A**, loading weights of different cytokines into the first principal component of the PCA in relation to their influence on depression or fatigue. A positive loading indicates that increased levels of these markers are related to increased depression or fatigue scores. **B & C**, relationship between first principal component and (B) depression score or (C) fatigue score. Linear regression line fitted with shaded area showing 95% confidence interval.

These data indicate that while no single marker was associated with depression or fatigue in our cohort, these data indicate that both symptoms are associated with a coordinated shift in the balance of cytokines and other blood markers. BMI had a modest influence on both fatigue and depression. Our data suggest that the effect of BMI and inflammation are additive in explaining the variability in depression score, whilst the influence of fatigue and inflammation could not be reliably disambiguated. Depression in our cohort was uniquely associated with increases in B-cell activating factor (BAFF) and CXCL1 and a decrease in PYY while fatigue was uniquely associated with increased IL1Ra, IL2Ra and CCL3 alongside a decrease in IL17D. Interestingly, both fatigue and depression were associated with increases in erythropoietin and leptin, and decreases in eotaxin / CCL11.

### Depression in psoriatic disease correlates with thalamostriatal connectivity in humans

We carried our resting state functional magnetic resonance imaging (rs-fMRI) on individuals with PsD before treatment with biologic anti-cytokine therapy and tested whether connectivity of individual brain regions with the whole brain correlated with the PROMIS depression score (figure 2A). We found that connectivity between the caudate and the right frontal pole significantly correlated with the depression score, but no significant correlation with subcallosal or orbitofrontal cortices (figure 2A). These individuals also scored highly on measures of impaired motivation and fatigue (supplementary table 1).

**Figure 2:**
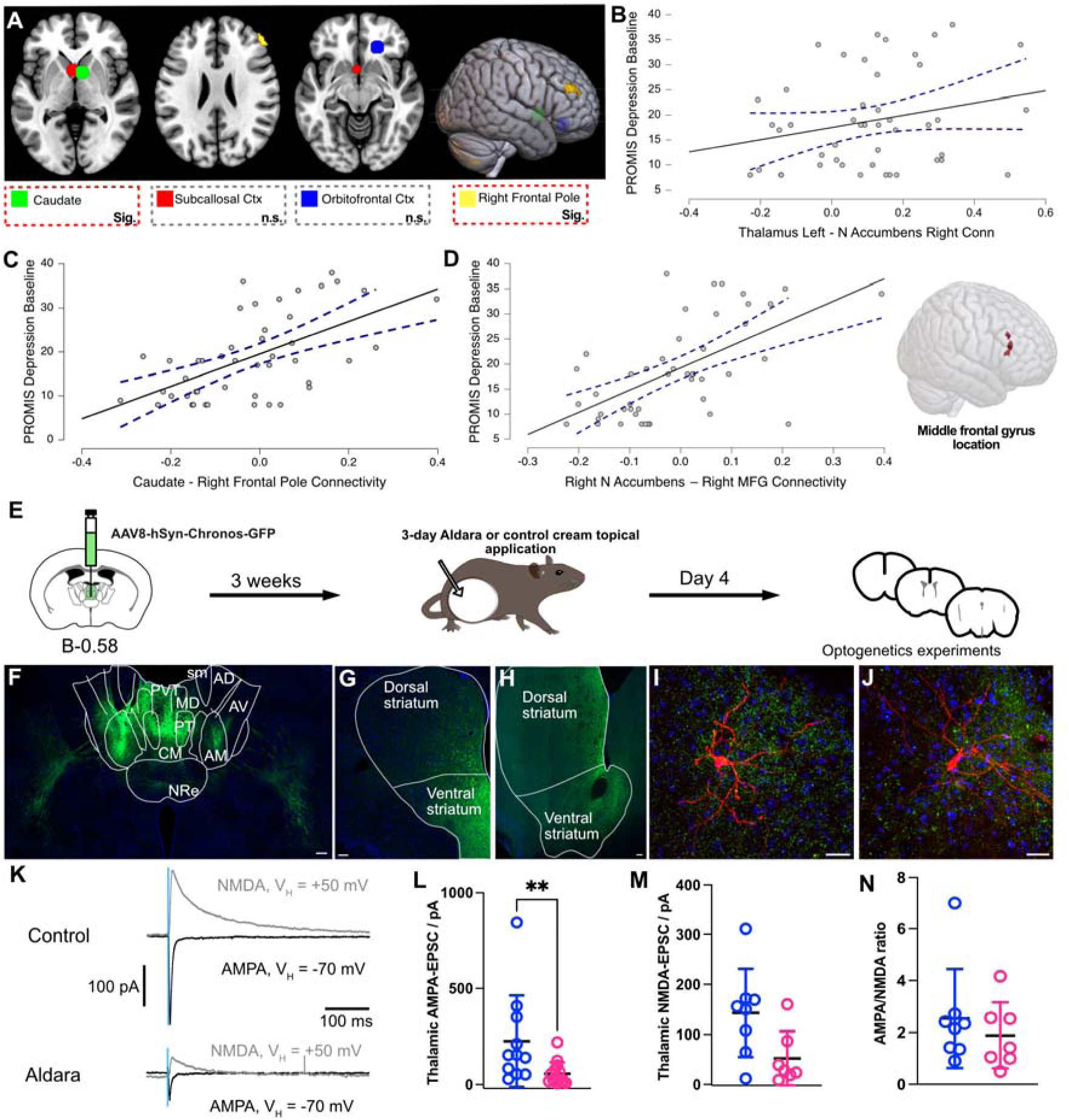
Psoriatic phenotype is associated with altered thalamostriatial signalling in both humans and mice. **A**, MNI template with the four regions-of-interest ROIs based on the Harvard-Oxford Atlas. We compared connectivity of each region with the whole brain and found that resting state MRI connectivity of caudate nucleus and right frontal pole, but not subcallosal or orbitofrontal cortices, correlated with the PROMIS depression score in patients with PsA. We also that the PROMIS depression score significantly correlated with connectivity between **B**, left thalamus and right nucleus accumbens (r = 0.33, FDR-corrected p=0.03) of patients with PsD (n = 46); **C**, caudate and right front pole (r = 0.58, FDR-corrected p=0.01) and **D**, right nucleus accumbens and right middle frontal gyrus (r = 0.64, FDR-corrected p=0.02). **E**, Schematic of experimental design for optogenetic and *ex vivo* patch clamp electrophysiology experiments investigating effect of Aldara treatment on midline thalamus projections to ventral MSNs. **F**, Representative image of midline thalamus showing expression of AAV8-hSyn-Chronos-GFP vector 3 weeks after injection. **G & H**, Expression of thalamic fibres in rostral and caudal portions of striatum, respectively. **I & J**, *Post hoc* recoveries of biocytin-filled MSNs in ventral striatum (red) with thalamic axons expressing Chronos-GFP (green); DAPI stain in blue. **K**, Representative traces of voltage-clamp electrophysiological recordings showing AMPA and NMDA receptor-mediated components of the optogenetics-evoked midline thalamic EPSC onto MSNs in the ventral striatum for control (top) and Aldara-treated (bottom) mice. **L**, Aldara treatment caused a significant reduction in the AMPA-receptor mediated thalamic EPSC, control (**i**) *vs* Aldara-treated (**ii**): 225.9 ± 71.7 pA *vs* 57.07 ± 15.9 pA, p = 0.0013, as measured with Mixed effects analysis; but **M**, had no statistically significant effect on the NMDA-receptor mediated component, control *vs* Aldara-treated: 142.9 ± 31.2 pA *vs* 52.29 ± 20.3 pA, p = 0.3486, as measured with mixed effects analysis. **N**, Aldara treatment had no significant effect on the AMPA/NMDA ratio of the thalamic input into ventral MSNs, control *vs* Aldara-treated: 2.54 ± 0.67 *vs* 1.89 ± 0.5, p = 0.6126, as measured with Mann-Whitney test. *Abbreviations: AD, anterodorsal thalamus; AM, anteromedial thalamus; AV, anteroventral thalamus; CM, centromedial thalamus; MD, mediodorsal thalamus; NRe, nucleus reuniens; PVT, paraventricular thalamic nucleus.* All data taken from at least 4 mice per group (across 3 different litters, with only 1 neuron taken per brain slice.

We next measured whether connections between pairs of brain regions displayed any significant correlation with the PROMIS depression score in our clinical population. In support of our main hypothesis, we found a positive correlation in functional connectivity between left thalamus and right nucleus accumbens with PROMIS depression score (figure 2B). Along with the thalamus and striatum, the prefrontal cortex is the third part of a brain circuit that has been implicated in depression^20^. Supporting with this, we also found significant correlations of functional connectivity between caudate and the right frontal pole (figure 2C), and right nucleus accumbens with the right middle frontal gyrus (figure 2D), with the PROMIS depression score. Additionally, we saw non-significant but trending-positive correlation between thalamic amplitude of low-frequency fluctuation (ALFF - a proxy measure of spontaneous neuronal activity^21^) and depression score (supplementary figure 2). Our clinical neuroimaging data align with a recent human neuroimaging study that found higher connectivity between paraventricular thalamic nucleus and ventral striatum in individuals with anhedonia^22^ Similarly, there was a positive correlation in functional connectivity between right nucleus accumbens and right middle frontal gyrus with fatigue, although this was mainly driven by the association of fatigue with depression (supplementary figure 2). We saw no significant correlation in functional connectivity between anterior cingulate cortex and either nucleus accumbens or thalamus with depression (supplementary figure 3).

### Psoriasis-like inflammation drives changes in thalamostriatal connectivity in mice

We next sought to determine whether mice displayed similar changes in thalamostriatal connectivity. We carried out stereotaxic surgeries to deliver AAV vectors to transduce expression of the excitatory opsin Chronos-GFP^23^ to neurons in midline thalamus, with an injection site targeting the border of the paraventricular thalamic nucleus (PVT) and the centromedial thalamus (CM, figure 2E). This led to widespread expression of Chronos-GFP in thalamic axons spread throughout ventral striatal regions including nucleus accumbens (figure 2F – I and supplementary figure 4). We made patch-clamp recordings in voltage-clamp mode from medium spiny neurons (MSNs) in ventral striatum, confirmed using *post hoc* morphological recovery (figure 2I & J). After 3 days of Aldara treatment, we found a significant reduction in the AMPA receptor-mediated component of the midline thalamic EPSC onto ventral MSNs in Aldara-treated mice relative to controls (figure 2K & L). However, we found no statistically significant reduction in the NMDA receptor-mediated component of the EPSC (figure 2K & M). While the statistical test did not reach the threshold for significance, the data appeared to be trending towards a reduction in Aldara treatment. Indeed, we saw no significant change in the ratio of AMPA/NMDA currents due to Aldara treatment, supporting the conclusion that both glutamatergic currents were likely reduced after Aldara treatment (figure 2N).

To test whether the neuroinflammation-driven change in thalamic EPSC may be due to altered presynaptic release, we examined spontaneous EPSCs of ventral MSNs (supplementary figure 5B). We found no significant effect of Aldara treatment on either the amplitude (supplementary figure 5C) or frequency (supplementary figure 5C) of spontaneous EPSCs. Finally, we found that treatment with Aldara did not significantly alter the input resistance of MSNs (input resistance, control (n=17) *vs* treated (n=26): 65.2 ± 6.3 MΩ *vs* 62.5 ± 6.2 MΩ; p = 0.4057, Mann-Whitney test), suggesting no change in intrinsic excitability of MSNs, consistent with a selective alteration of thalamostriatal synapses. Furthrmore, a secondary analysis of our whole brain RNAseq transcriptomic dataset^24^ revealed a significant downregulation in genes relevant to glutamatergic transmission (Homer1, Homer2, Glul and Gls) as well as astrocyte-specific EAAT2 implying an impairment in astrocytic glutamate reuptake (supplementary figure 5A).

### Impaired motivation and anhedonia-like behaviours in mice modelling psoriatic disease

After confirming that perturbations to thalamostriatal signalling occurred both in humans with PsD and Aldara-treated mice modelling psoriasis, we next tested whether the murine model displayed similar impairments in motivational behaviour. We carried out behavioural assays after topical application of Aldara cream for consecutive 3 days, measuring in-cage behaviours daily and other tests on the day after the final application. We used the nest-building assay^25^ as a measure of intrinsic motivation in mice. Mice were provided with 3.0 g nestlet each evening and we measured the amount of unused nestlet the following morning. After 3 days of treatment, we observed a significant increase in the amount of unused nestlet in Aldara-treated mice relative to controls (figure 3A). Additionally, the quality rating of the nest built by the mice over the 24h period progressively decreased in Aldara-treated mice, reaching significance at day 3 post-treatment (figure 3B). In contrast, control mice displayed a trend for a progressive improvement in nest quality. Examples of nests, corresponding to each of the possible grades for nest-building assay, are shown in supplementary figure 6. Overall, this test suggested a reduction in intrinsic motivation in Aldara-treated mice. To further assay anhedonia-like behaviour, we used the sucrose preference test. We measured the sucrose-to-water ratio (weight change in sucrose solution divided by the weight change in water solution in the previous 12 hours) twice daily, to account for the nocturnal cycle of mice: their baseline preference for sucrose was absent during the daytime (typical ratio of 1 for both groups) but was on average ∼4 during the night in both groups. Aldara treatment significantly reduced the preference for sucrose at night, relative to control-treated mice (figure 3C). Together, the nest-building assay and sucrose preference test indicate an emerging anhedonia-like phenotype after 3 days treatment with Aldara.

**Figure 3:**
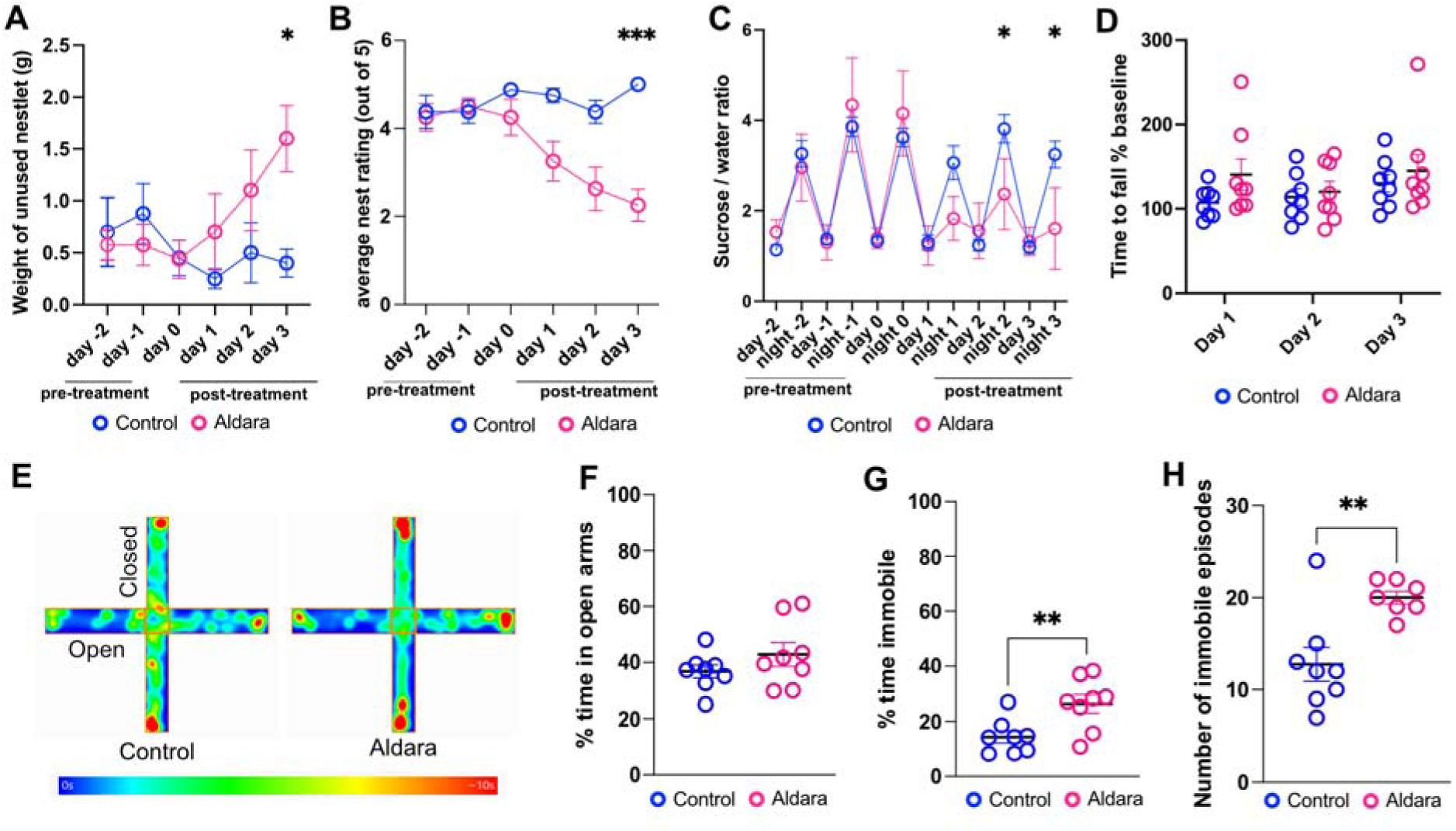
Behavioural assessment of 3-day Aldara-treated mice. In all panels control mice are shown in blue and Aldara-treated mice in pink. **A**, Nest-building assay across treatment time: significant interaction between Aldara treatment with time on amount of unused nestlet (p = 0.0079 in 2-way repeated measures ANOVA), with a significantly more unused nestlet in Aldara-treated mice by day 3 (control *vs* Aldara-treated 1.1 ± 0.2 g *vs* 0.38 ± 0.1 g, p = 0.0403 in *post hoc* Šídák’s multiple comparisons). **B**, Aldara treatment caused increasingly lower nest-building ratings across time which reached significance on day 3 (Effect of treatment: p = 0.0007; effect of time: 0.0067; interaction: p < 0.0001; two-way ANOVA with Šídák’s *post hoc* tests; 5 is highest and 1 is lowest). **C**, Sucrose preference at day and night during treatment. Significant interaction between treatment and time (p < 0.0001; two-way ANOVA) with Aldara-treated mice showing significantly less preference for sucrose during the night after 2 and 3 days of treatment (Šídák’s *post hoc* tests). **D**, Rotarod performance (% time to fall from the rod compared to baseline) was evaluated every day prior to cream treatment, control *vs* Aldara-treated 117.1 ± 11.73 vs 135.1 ± 13.2 % show no significant change as found with a two-way ANOVA. **E**, Representative heatmaps from elevated plus maze (EPM) recordings performed 24h following final treatment. **F**, Percentage time spent in open arms of the EPM was not changed, p = 0.2276, unpaired two-tailed t-test. **G**, Percentage time spent immobile (immobility defined as not moving for longer than 2 seconds) in 5-minute trial, control *vs* Aldara-treated p = 0.0093, unpaired two-tailed t-test. **H**, Total number of immobile episodes during EPM testing, p = 0.0039, unpaired two-tailed t-test. Scatter plots display data points (each point is one animal) and mean ± SEM; n=8 mice per group in all tests; *, p < 0.05; **p < 0.01; *** p < 0.005.

We used the rotarod test to assess motor capability in evoked movement in Aldara-treated mice. There was no significant difference in rotarod performance (time to fall relative to baseline) between treatment groups (figure 3D), demonstrating there were no motor impairment due to Aldara treatment. In a separate cohort of mice, we carried out the elevated plus maze test on the day after the final Aldara treatment (figure 3E). We saw no difference in preference for the closed and open arms in Aldara-treated mice relative to controls (figure 3F). However, we found that Aldara-treated mice spent a significantly longer period immobile (figure 3G) relative to control mice. Additionally, these mice exhibited a significantly greater number of immobile periods (figure 3H) on the elevated plus maze than control mice, demonstrating a start-stop like activity within the test. We concluded that the reduced mobility seen in behavioural testing was likely to reflect reduced motivation to carry out exploratory behaviour and not impairments in the ability to move due to treatment.

### Neuroinflammation changes neuronal activation in a region-specific manner

Having confirmed that mice modelling psoriasis showed disruptions to the same thalamostriatal circuitry as humans and display similar anhedonia / depression-like behaviours, we next sought to determine whether the neurobiological changes in mouse displayed the same regional specificity as observed in humans with PsD (figure 2A). Using a spatial transcriptomics approach, we previously reported that neuroinflammation in the Aldara model is associated with global increases in expression genes associated with inflammation and glial reactivity, alongside regionally-specific changes in expression of genes related to neurotransmission^24^. Here, we tested whether these transcription-level changes were reflected at the protein level. First, we first assayed neural activity using immunohistochemistry for the immediate early gene product c-Fos, an established marker of neuronal activity. After 3 days of Aldara treatment, when behavioural changes are apparent, we saw a reduction in synaptic strength of thalamic input to ventral striatum (figure 2K). We previously shown via mass spectrometry that Aldara enters the brain within 4 h of treatment^26^, so measured c-Fos reactivity at this point to determine which brain regions are first affected. We saw a significant increase in c-Fos activity within the midline thalamus of Aldara-treated mice relative to controls (supplementary figure 7B), indicating that midline thalamus is engaged at the earliest stage of neuroinflammation. We next used anatomical methods to quantify glutamate transporters in ventral striatum after 3 days of Aldara treatment, when behavioural effects were most pronounced. Ventral striatum receives input from both prefrontal cortex and midline thalamus, so we used immunofluorescence to visualise glutamate transporters VGlut1 and VGlut2, which are preferentially expressed on cortical and thalamic axons, respectively^27^, to parse these two striatal inputs (supplementary figure 7C). We chose vS1 as a control region due to the strong thalamic innervation of layer 4 in the barrel cortex (supplementary figure 7E). We found a significant increase in the number (supplementary figure 7D) and size (supplementary figure 7E) of VGlut2 axon terminals in ventral striatum but not vS1 and observed no significant changes in VGlut1-containing axon terminals. These data suggest that there was a change in the thalamic, but not cortical, inputs to ventral striatum and the lack of change in vS1 suggests that this was specific to the reward pathway.

### Neuroinflammation drives functional changes in microglia and astrocytes

Having established that psoriasis-like neuroinflammation drives was associated with changes in thalamostriatal function related to anhedonia or depressive-like symptoms across species, we used the mouse model to further probe the underlying mechanisms of neuroinflammation. While microglia have long been seen as the ‘immune cells’ of the brain, astrocytes also play a key role as both sensors and effectors of the immune system^28^. We employed flow cytometry and immunohistochemistry to characterise functional changes in astrocytes and microglia in the Aldara model. IBA-1 immunohistochemistry experiments revealed changes in microglial morphology (less ramified, more amoeboid) indicative of a reactive state (figure 4A). Despite these obvious qualitative changes in morphology, we see no significant quantitative changes in the total area covered by IBA-1 (figure 4B), with hippocampus being the only region showing a significant increase relative to control mice. Flow cytometry gating with CD45^int^ and CD11b^hi^ for microglia showed increase in the total number of microglia in Aldara treated compared to controls (figure 4C). We saw significant increase in microglia expressing markers of reactivity, MSR1, F4/80 and MHCII (figure 4D), confirming that Aldara treatment drives microglia into a reactive phenotype. Further analysis of the whole-brain transcriptome of our previously reported data^24^ confirmed that several genes associated with microglial activation were upregulated in Aldara-treated mice (supplementary figure 8A).

**Figure 4:**
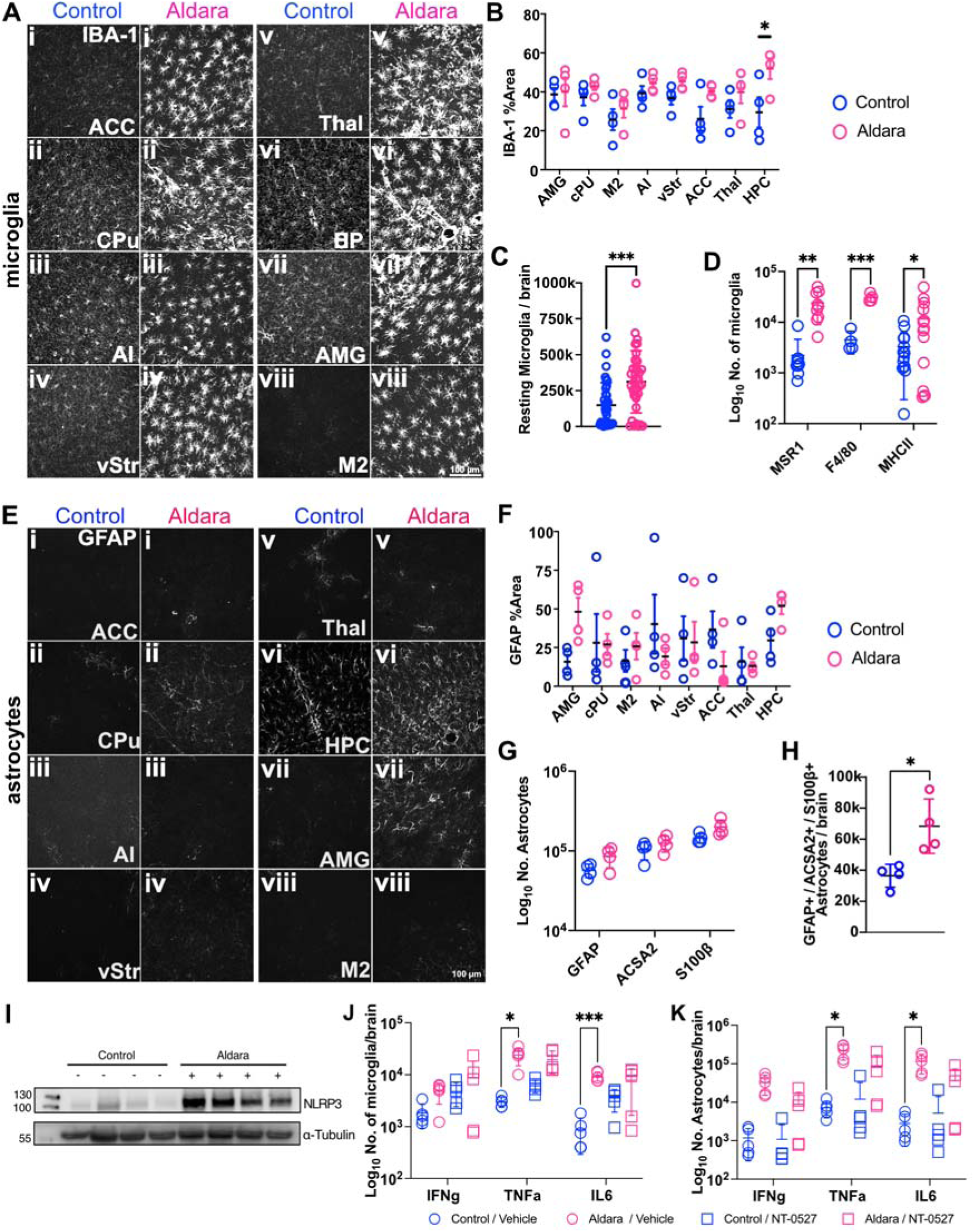
Aldara treatment induces reactivity in microglia and astrocytes driven via mechanisms involving NLPR3 inflammasome. In all panels control mice are shown in blue and Aldara-treated mice in pink. **A**, images taken from immunohistochemistry stain for IBA1 in control- and Aldara-treated brains for: **i**, anterior cingulate cortex (ACC); **ii**, caudate putamen/dorsal striatum (CPu); **iii**, agranular insular cortex (AI); **iv**, nucleus accumbens/ventral striatum (vStr); **v**, midline thalamus (thal); **vi**, dorsal hippocampus (HPC); **vii**, amygdala (AMG); **viii**, motor cortex (M2). **B**, total % area of section labelled for IBA1. **C & D**, flow cytometry counts for total number of cells in brain homogenate from control-and Aldara-treated mice, selecting for the following microglia markers: total resting microglia (CD45int and CD11bhi; (Control *vs* Aldara 165242 ± 158635 *vs* 313437 ±229741 p < 0.0001, two-tailed t-test. **D**), MSR1, F4/80 and MCHII 2221 ± 2391 *vs* 24098 ± 15110 for MSR1, p = 0.0024, 4499 ± 2130 *vs* 30549 ± 5389 for F4/80, p = 0.0009, 3283 ± 2985 *vs* 11665 ± 14100, p = 0.0470, in *post hoc* Šídák’s multiple comparisons test after two-way ANOVA. **E**, images taken from immunohistochemistry stain for GFAP in control- and Aldara-treated brains for same regions as **A**. **F**, total % area of section labelled for GFAP for same samples. **G**, flow cytometry counts for total number of cells in brain homogenate from control- and Aldara-treated mice, positive for GFAP, ACSA2 and S100β astrocytic markers. 57040 ± 10912 *vs* 85302 ± 24515 (GFAP), 103243 ± 25997 *vs* 124847 ± 28099 (ACSA2), 144611 ± 19643 *vs* 198835 ± 43995, no significant changes as found in *post hoc* Šídák’s multiple comparisons test after two-way ANOVA. **H**, Aldara treatment significantly increased the number of astrocytes that were triple-labelled for GFAP, ACSA2 and S100β. **I**, Western blot shows an increase in the NLRP3 in the brain **in** Aldara treated group as compared to control. **J**, Flow cytometry Number of microglia, expressing IL-6, but not TNFα and IFN-γ, is reduced in Aldara-treated animals treated with NLRP3 antagonist, as seen with flow cytometry, n=5. ***TNF***α: control / vehicle: 2998 ± 649; Aldara / Vehicle, 23931 ± 9029; control / NT-0527, 6163 ± 1689; Aldara / NT-0527, 19534 ± 9074. ***IL-6***: Control / Vehicle,875 ± 582; Aldara / Vehicle, 9397 ± 1876; control / NT-0527, 3550 ± 1658; Aldara / NT-0527, 3550 ± 1658. ***IFN-***γ: control / vehicle, 1828 ± 897; Aldara / vehicle, 4780 ± 2067; control / NT-0527 4740 ± 2320; Aldara / NT-0527, 8859 ± 9317. All statistical comparisons *vs* control / vehicle; *post hoc* Šídák’s multiple comparisons test after two-way ANOVA. **K**, Number of astrocytes, expressing IFN-γ, TNFα and IL-6, is reduced in Aldara-treated animals treated with NLRP3 antagonist, as seen with flow cytometry, n=5. ***TNF***α: control / vehicle, 7245 ± 3343; Aldara / vehicle, 215313 ± 90628; control / NT-0527, 12075 ± 20288; Aldara / NT-0527, 80809 ± 8290. ***IL-6***: control / vehicle, 2803 ± 2079; Aldara / vehicle 114687 ± 60531; control / NT-0527, 5205 ± 8883; Aldara / NT-0527, 36195 ± 37113. ***IFN-***γ: control / vehicle, 1181 ± 882; Aldara / vehicle, 32601 ± 17405; control / NT-0527, 1083 ± 1521; Aldara / NT-0527, 8859 ± 9317 p=0.3698. Panels **J & K**: comparisons *vs* control / vehicle; *post hoc* Šídák’s multiple comparisons test after two-way ANOVA. *, p <0 .05; ***, p < 0.005.

We carried out the same analysis in astrocytes, using GFAP as a morphological marker for immunohistochemistry (figure 4E). Unexpectedly, we failed to detect major changes in astrocyte immunoreactivity or morphology in response to Aldara treatment, nor were there changes in average area covered by GFAP (figure 4F). GFAP, S100β and ACSA2 (figure 4G) were used as markers to identify potential astrocytic activation in flow cytometry, but we found little evidence of differences in the individual expression profile of these three markers in treated brains compared to control. However, there was a significant increase in the number of triple-positive astrocytes for GFAP, S100β and ACSA2 (figure 4H) in the Aldara treated mouse brains, which could provide putative evidence of reactivity. Transcriptomic analysis also found upregulation of genes associated with astrocytic activation in Aldara-treated mice (supplementary figure 8B). In summary, this experiment found strong evidence of microglial reactivity and weak evidence suggesting astrocytic reactivity in response to Aldara treatment.

Reactive astrocytes and microglia both produce cytokines that can act as a positive feedback mechanism to drive further inflammation. Astrocytic reactivity in inflammatory states is driven by reactive microglia^29^, a process that depends upon activation of the NLRP3 inflammasome in microglia^30–32^, signalling via IL-1β. We found that treatment with Aldara significantly increased NLRP3 protein expression in mouse brain (figure 4I). To test our dual hypotheses that both glial types increased cytokine production in the Aldara model and that inflammasome is mediating this, we employed a 2 x 2 experimental design to measure cytokine production in glial cells using flow cytometry. We employed a recently described brain-penetrant inflammasome inhibitor, NT-0527^33^, in an attempt to modulate the neuroinflammatory response to Aldara treatment. Mice were pre-treated with NT-0527 at 100 mg/kg orally (or vehicle) for 24h before Aldara treatment and then once daily during the Aldara protocol. We confirmed that both microglia (figure 4J) and astrocytes (figure 4K) increased cytokine production in response to Aldara treatment, and that the NLRP3 inflammasome inhibitor could indeed significantly attenuate cytokine production in both cell types. Treatment with NT-0527 also blocked Aldara-driven increase in circulating monocytes and neutrophils found in the blood (supplementary figure 9), confirming that this compound had both central and peripheral effects. This experiment confirmed that both astrocytes and microglia became reactive by increasing production of TNF and IL-6 in the Aldara-driven psoriasis model, and that this positive-feedback mechanism is at least in part mediated via the NLRP3 inflammasome.

### Immune cell infiltration into the brain following Aldara treatment

We have previously shown that 3 days of treatment with Aldara results in development of psoriasis in mice alongside an ingress of peripheral immune cells into the brain^34^, corresponding to when we observed synaptic and behavioural changes. Multiple peripheral immune cell types, including T cells and monocytes, can produce cytokines and are capable of driving microglial and astrocytic reactivity. Consequently, we used flow cytometry (see supplementary material for gating strategy) and immunohistochemistry experiments to determine which immune cell type(s) were entering the brain in a response to inflammation. Flow cytometry of brain single-cell homogenate showed expression of 4 key populations based on two classical makers CD45 and CD11b (figure 5A). The four cell populations were: non-immune (CD45^-^CD11b^-^); lymphoid (CD45^+^CD11b^-^); myeloid (CD45^hi^CD11b^+^); and microglia (CD45^int^CD11b^+^)^35^. Aldara treatment caused a substantial increase in all immune cell types in brain homogenates (figure 5A) with a significant overall increase in the total number of live cells (figure 5B) relative to controls. This increase was attributable to CD45^+^, indicating a 4-fold increase in the number of immune cells (including microglia) in the brain (figure 5C). Of the peripheral immune cells infiltrating the brain, this population was primarily comprised of monocytes, B cells, NK, and NKT cells (figure 4D; note that data are plotted on a logarithmic scale). We also saw a substantial ingress of CD3^+^ T cells into the brain following Aldara treatment (figure 5E) and, of these, we saw significantly increased level of both CD4^+^ and CD8^+^ T cells (figure 5E – F). Regionality of infiltrating T cells was assessed using immunohistochemistry. We found a uniform increase in CD4^+^ cells in all brain regions tested (figure 5G – 5H), although the density of CD4^+^ T cells did not reach the significance threshold on *post hoc* multiple comparisons tests for hippocampus and ventral striatum (figure 5H) despite their presence in these regions. Thus, both flow cytometry and immunohistochemistry confirm that Aldara treatment is associated with substantial peripheral immune cell ingress into the brain, with an apparently uniform brain parenchymal penetration.

**Figure 5:**
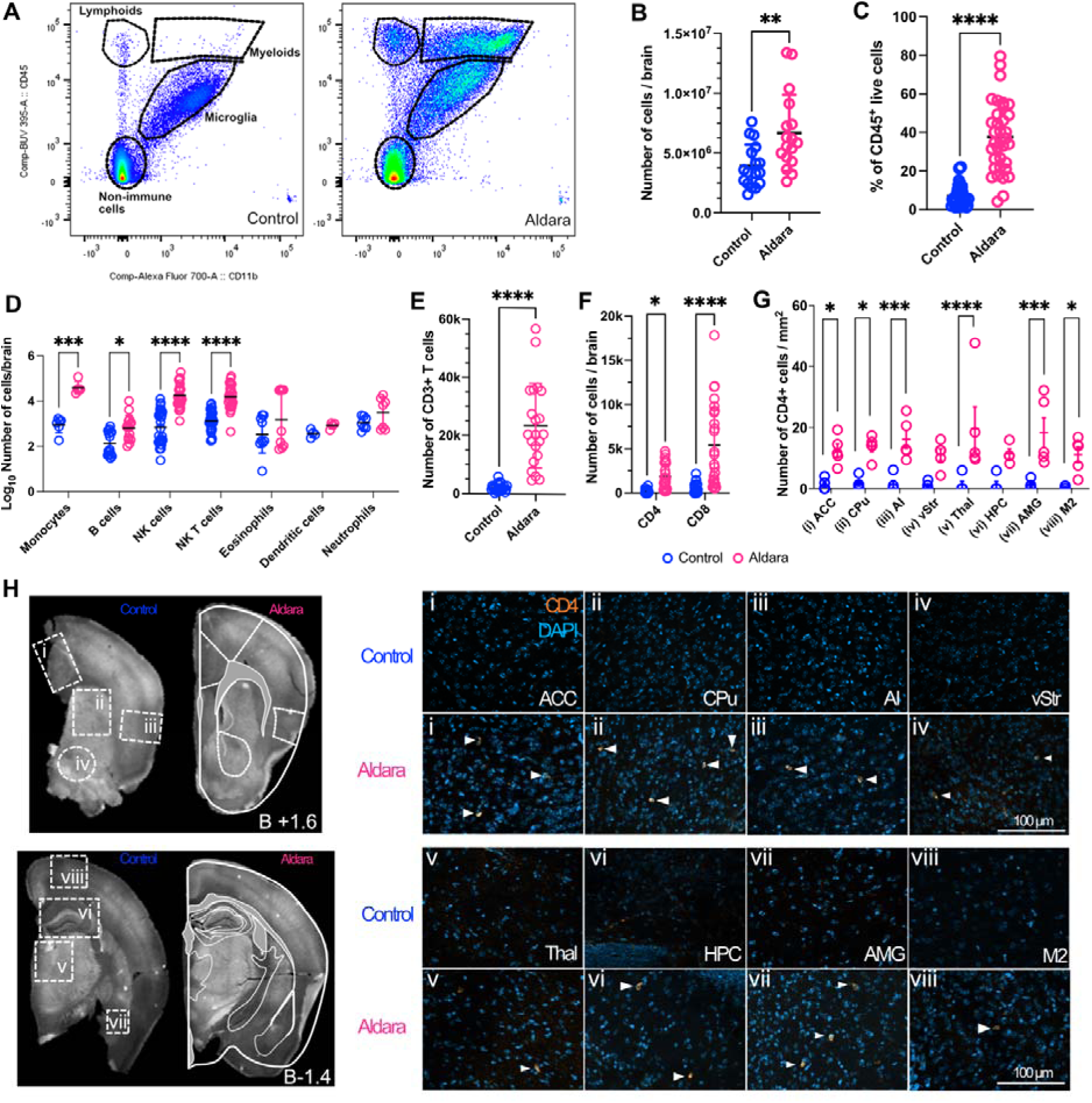
Ingress of peripheral immune cells into the brain is observed after 3 days of Aldara treatment. In all panels, control mice are shown in blue and Aldara-treated mice in pink. **A**, Representative basic flow cytometry panels from a single control (left) and Aldara-treated (right) brain. The 4 broad cell types are indicated within dashed lines. **B**, total number of cells (10^^6^) found in the brain homogenates from control and Aldara-treated mice, n=17 4± 2 *vs* 7 + 3 p = 0.0039, unpaired t-test. **C**, relative proportion of live CD45+ cells in control- and Aldara-treated brain homogenates control *vs* treated 38 ± 3% *vs* 7 ± 1%, n=36 mice per group; p < 0.0001, unpaired t- test. **D**, numbers of different immune cell types, as measured with flow cytometry, in control- and Aldara-treated brain homogenates (data plotted on log scale) Monocytes n=6 1061 ± 533 *vs* 47317 ± 40268 p < 0.0001; B cells n= 15 248 ± 245 *vs* 1510 ± 2593 p = 0.0170; NK cells n= 27 2172 ± 2995 *vs* 34584 ± 44310 p < 0.0001; NK T cells n= 27 2068 ± 2030 *vs* 28573 ± 31534 p < 0.0001; Eosinophils n= 9 9257 ± 1021 *vs* 13923 ± 16315 p = 0.1606; Dendritic cells n= 4 349 ± 148 *vs* 871 ± 255 p = 0.9825; Neutrophils n= 7 1295 *±* 791 *vs* 7016 ± 7079 p = 0.7446 as shown by Sidak’s multiple comparison test. **E**, absolute numbers of CD3-expressing T-cells in control and Aldara-treated mice n=21 control *vs* treated 2150± 1320 *vs* 23579 ± 14547 p < 0.0001 unpaired t- test. **F**, Aldara treatment led to a significant increase in the numbers (10^^3^) of CD4 and CD8 expressing T –cells CD4 T cells, n=24 control vs treated 188 ± 192 *vs* 1850 ± 1461 p = 0.0245 and CD8 T cells control vs treated 478 ± 563 *vs* 5402 ± 4781 p < 0.0001 as shown by two-way ANOVA. **G**, Summary immunohistochemistry data for density of CD4+ T cells in multiple brain regions from control versus Aldara-treated brain sections: **i**, anterior cingulate cortex (ACC) n=5 1.2 ± 1.8 *vs* 12.5 ± 4.7 p<0.05; **ii**, caudate putamen/dorsal striatum (CPu) n=5 1.6 ± 1.97 *vs* 13.7 ± 3.6 p<0.05; **iii**, agranular insular cortex (AI) n=5 1.4 ± 2.7 *vs* 16.1 ± 6.7 p<0.001; **iv**, nucleus accumbens/ventral striatum (vStr) n=5 0.8 ± 1.2 *vs* 11.1 ± 4.4 p=0.0608; **v**, midline thalamus (thal) n=5 1.2 ± 2.6 *vs* 19.6 ± 16.1 p<0.001; **vi**, dorsal hippocampus (HPC) n=5 1.2 ± 2.6 *vs* 11.4 ± 3.3 p=0.0965; **vii**, amygdala (AMG) n=5 1.1 ± 1.5 *vs* 18.3 ± 11.1 p<0.001; **viii**, motor cortex (M2) n=5 0.4 ± 0.5 *vs* 11.1 ± 5.7 p<0.05. **H**, Representative example of coronal slices taken from anterior (Bregma +1.6, left) and posterior (Bregma −1.4, right) brains. Brain regions for CD4^+^ quantification are highlighted. Representative images of each region from control and Aldara-treated brain sections are shown (CD4 orange, DAPI blue), positive cells are indicated by white arrowheads. Scale = 100 µm. Each point represents an individual animal (4 animals per experiment and cumulative at least 4 – 5 experiments per flow result), mean ± S.E.M. Unpaired t-tests (with or without Welsch’s correction as appropriate) were carried out on data from B, C, E, F; two-way ANOVAs were carried out on data on D and G. *, p < 0.05; **, p < 0.01; ***, p < 0.005; ****, p < 0.0001.

To further define the T cell population entering the brain, we carried out a subsequent experiment and found significant increases in subsets of infiltrating CD4^+^ and CD8^+^ T cells in brain of treated mice relative to controls, with increases in Th1/Tc1 cells secreting TNFα, but with no Th2 cells secreting the anti-inflammatory IL-4 (supplementary figure 10A & D, respectively). These data indicate a Th1/Tc1 signature of T cells in brain; there was some evidence of a Th17/Tc17 response, but the increase in CD4^+^IL-17^+^ and CD8^+^IL-17^+^ cells in Aldara-treated mice did not reach significance, likely due to high variability in the data. In this second experiment, we also used flow cytometry to internally replicate our finding that both microglia and astrocytes increased expression of proinflammatory cytokines (TNFα, IL-6 and IFN-γ). Aldara treatment significantly increased the number of microglia producing all 3 cytokines relative to controls (supplementary figure 10E), and the number of astrocytes producing IL-6 and IFN-γ (supplementary figure 10F). We also observed a significantly increased proinflammatory signature in infiltrating monocytes, which produced both TNFα and IL-6 in Aldara-treated mice (supplementary figure 10G). Further analysis of the whole brain transcriptome dataset revealed upregulation of genes for proinflammatory cytokines and chemokines in Aldara-treated mice (supplementary figure 10H). These chemokines are key to immune cell ingress to brain.^36^

### Contribution of different peripheral immune cell populations to neuroinflammation

Finally, we aimed to determine the contributions made by different types of infiltrating immune cells to our observed phenotypes in the mouse psoriasis model, focusing on monocytes and T cells. We used systemic treatment of antibodies against CD4 and CD8 to deplete circulating T cell populations to limit their entry to the brain. We used iCCR ^-/-^mice lacking the chemokine receptors CCR1, CC2, CCR3 and CCR5^37^ to inhibit egress of monocytes from bone marrow into the periphery and their subsequent ingress to the brain. Our control experiments using flow cytometry validated the approach: following Aldara treatment, T cell depletion resulted in a substantial knock-down of T cells in blood, brain and spleen (supplementary figure 11), and in iCCR ^-/-^mice there was a substantial reduction in circulating monocytes (supplementary figure 12).

Depletion of CD4+ and CD8+ T cells did not substantially alter the profile of the immune cells seen with Aldara treatment, with the main populations of myeloid, lymphoid and microglia apparent (figure 5A). While we substantially reduced the number of T cells entering the brain, we still observed significant ingress of other leukocytes (NK, NKT and B cells) and myeloid cells such as microglia and macrophages (supplementary figure 12A). Aldara treatment on the iCCR^-/-^mice did show a reduction in the myeloid population after Aldara treatment (figure 6A), as expected, representing the substantial decrease in the monocyte population. Lymphocytes such as T and B cells continued to infiltrate the brain after Aldara treatment in the iCCR^-/-^mice (supplementary figure 12B). Using weight loss and nest-building assays as proxy measures for Aldara-driven deficits in motivational behaviour, we found that neither reducing infiltrating monocytes in iCCR^-/-^mice (figure 6B & C) nor blocking T cell ingress (figure 6E & F) was sufficient to ameliorate these deficits. In the iCCR^-/-^ mice, but not in the T cell depletion mice, we did see a significant impairment in the Aldara-driven increase in numbers of microglia (or microglia-like) cells (figure 6D and G, respectively). We conclude that neither T cell nor monocyte ingress into the brain is individually necessary to drive the neuroinflammatory response seen in the Aldara model of psoriasis and, indeed, we observed morphological changes in microglia consistent with a reactive phenotype in the iCCRK^-/-^ mouse model (figure 6H).

**Figure 6:**
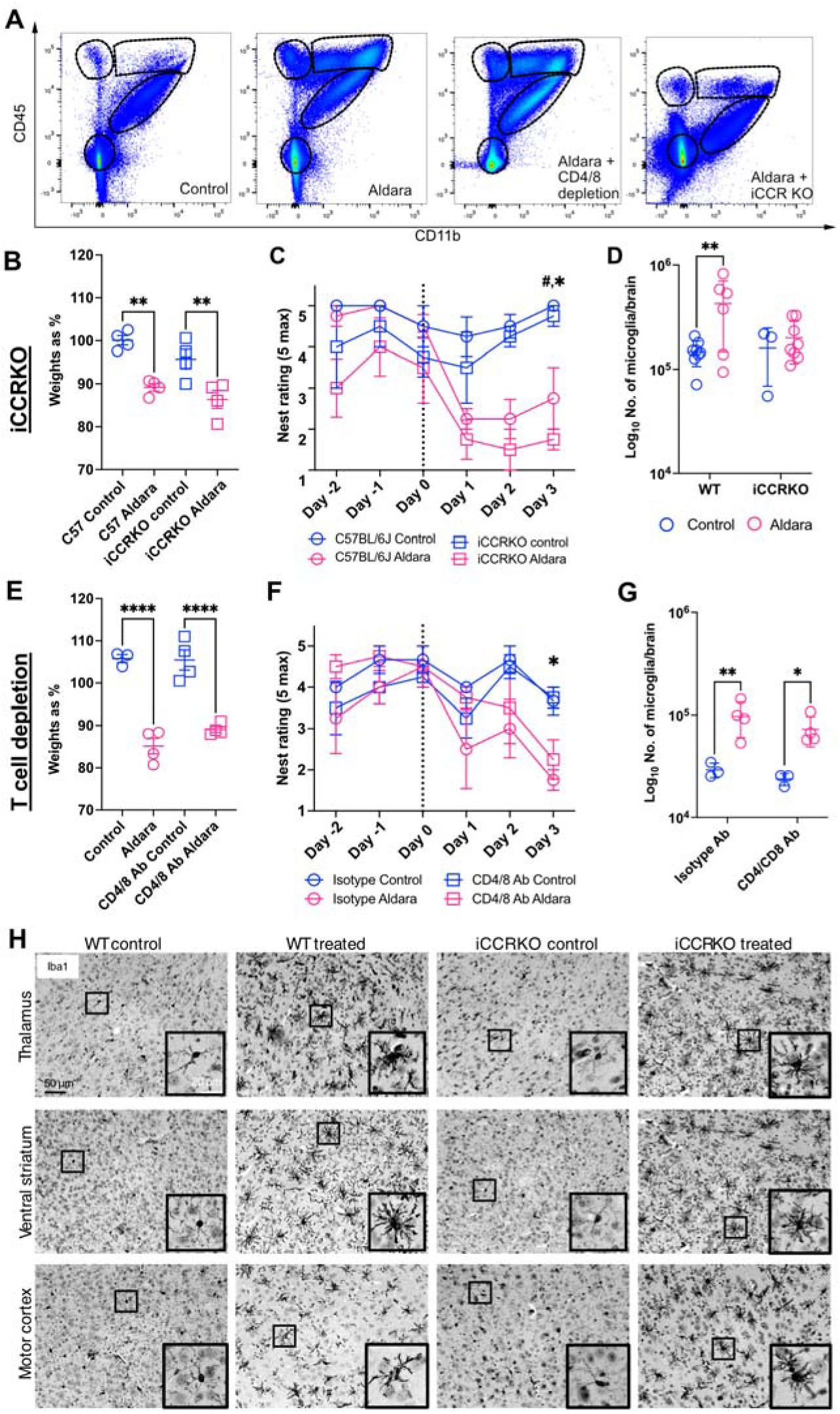
Neuroinflammatory phenotypes persist when immune cell ingress into the brain is reduced. **A**, Representative basic flow cytometry panels from WT control-treated, WT Aldara-treated, WT CD4/8 depleted control-treated, and iCCR ^-/-^ Aldara-treated mice. The 4 broad cell types are indicated within dashed lines. **B**, iCCRK ^-/-^ mice experience similar weight loss to wildtype mice during the 3-day Aldara treatment, C57BL6/J control *vs* treated n= 4, 100 ± 2 *vs* 89 ± 2 %, p = 0.0013, iCCR ^-/-^ control *vs* treated n= 4, 96 ± 5 *vs* 86 ± 4 %, p = 0.0039, as found with *post hoc* Šídák’s multiple comparisons test after one-way ANOVA. **C**, iCCR ^-/-^ and wildtype mice show similar impairments in nest building during the 3-day Aldara treatment. C57BL6/J control *vs* treated on cull day n=4 5 ± 0 *vs* 3 ± 2, p = 0.0133, iCCR ^-/--/-^ ontrol *vs* treated on cull day 5 ± 1 *vs* 2 ± 1, p = 0.0005, as found with *post hoc* Šídák’s multiple comparisons test after two-way ANOVA. **D**, iCCRKO mice show a smaller increase in the number of microglial cells in response to Aldara treatement relative to wild-type mice. **E**, CD4/8 depleted mice show a similar weight loss to control mice during the 3-day treatment. Isotype control *vs* treated, n=4, 98 ± 3 *vs* 84 ± 3 %, p < 0.0001, CD4/CD8 Ab control *vs* treated 96 ± 3 *vs* 80 ± 4 %, p < 0.0001, as found with *post hoc* Šídák’s multiple comparisons test after one-way ANOVA. **F**, CD4/8 depleted mice show similar impairments in nest building during the 3-day Aldara treatment. Isotype control *vs* treated, n=4, 4 ± 1 *vs* 2 ± 1, p = 0.0013, CD4/CD8 Ab control *vs* treated 4 + 1 *vs* 2+ 1, as found with *post hoc* Šídák’s multiple comparisons test after two-way ANOVA. **G**, CD4/8 depleted mice show a similar increase in the number of microglial cells as compared to wild-types Isotype control vs treated, n=4, 29199 ± 4806 *vs* 96917 ± 36607 p = 0.0182, CD4/CD8 Ab control *vs* treated 23738 ± 3220 *vs* 72612 ± 23573 p = 0.00927 as found with Tukey’s multiple comparison test after two-way ANOVA. **H**, representative images taken of IBA1-staining in brain sections from thalamus, ventral striatum and motor cortex following 3 days of control or Aldara-treatment in wild type and iCCR ^-/-^ mice. Aldara-treated iCCR ^-/-^ mice show a similar degree of microglial reactivity.

## Discussion

The aim of our study was to test the hypothesis that neuroinflammation plays a causal role in anhedonic phenotypes occurring in psoriatic disease by perturbing thalamostriatal circuity. In humans with PsD, we saw a positive correlation between depression scores and functional connectivity between thalamus, nucleus accumbens and prefrontal areas. We used a reverse-translation approach with Aldara to generate a psoriasis-like condition in mice and found that this model recapitulated the clinical features of altered thalamostriatal signalling and impaired motivational behaviours. The mouse model allowed us to probe neurobiological and immune mechanisms more deeply, with multiple lines of evidence converging upon reactive microglia and astrocytes mediating the neuroimmune response, with contributions from infiltrating peripheral immune cells.

Our working model is that inflammation quickly drives activity in the midline thalamus that, over time, leads to a reduction in efficacy of thalamostriatal signalling, likely due to changes in glutamate signalling through receptor turnover and impaired ability of astrocytes to regulate neurotransmitter levels. We found that the astrocytic cytokine production and reactivity is driven by microglia via NLPR3-dependent mechanisms, and that multiple populations of immune cells are capable of infiltrating the brain and exacerbating ongoing neuroinflammation.

### Relationship between inflammation and depression in humans with PsD

We found that depression in humans was associated a changes in a small number of markers, with BAFF in particular having a large influence on the depression score. BAFF, B-cell activating factor, is a member of the TNF receptor superfamily^38^. BAFF has not been widely studied in the context of depression, although a recent article found that increases in BAFF in haemodialysis patients predicted future development of depression^39^. Elevated plasma BAFF levels have also recently been implicated in schizophrenia and bipolar disorder^40^; we observed B-cell ingress into the CNS in our Aldara-treated mice. In another recent preclinical study it was shown that B-cells may influence stress-related behaviour by regulating meningeal myeloid cells activation^41^ CXCL1 was also positively associated with depression in our cohort. In mice, increases in CXCL1 has been shown to increase depressive-like behaviours in models of chronic unpredictable stress^42^ but *reductions* in CXCL1 in plasma have been associated with elderly depression in humans^43^. These contradictory results highlight the risk in studying single molecular markers in isolation and support our finding that the inflammation-driven depression is rather associated with disruptions within networks of immune-mediating molecules. It should be noted that our sample was relatively small (n=45) for a heterogenous population and we used stringent inclusion criteria for cytokines (excluding key molecules such as TNFa and IL6 due to these analytes being outwith the limits of detection for a substantial for a substantial subset of our participants). Despite this limitation, we still observed a significant, moderate effect size in our model.

Elevated fatigue was associated with increases in IL1Ra, IL2Ra and CCL3. The increases in IL1Ra are indicative of an anti-inflammatory response to increased proinflammatory IL1α and IL1β. IL2 is central to controlling regulatory T cells activity (T_reg_). The emerging view is that T_reg_ cells in the brain contribute to the attenuation of glial cell reactivity^44^. The upregulation of the antagonist receptor for IL2, which promotes T_reg_ activity, suggest an abnormality in this control mechanism that might lead to continuation of neuroinflammation. The effects of the increase in the myeloid cell-attracting chemokine CCL3 are seen in the myeloid cell ingress the brain in our preclinical data, again this could be in response to ongoing neuroinflammation or represent another potential source of proinflammatory cytokines within the brain. Increases in erythropoietin were associated with both depression and fatigue in our model. Interestingly, single doses of erythropoietin were found to alleviate depressive symptoms in individuals with MDD or bipolar disorder^45^, while resistance to erythropoietin is associated with fatigue in haemodialysis patients^46^. Our cohort of individuals are those living with PsD, which is a chronic inflammatory disease, so these data highlight the need to consider chronicity when looking for biomarkers of inflammation-driven depression. Although not one of the major components of the principal components in our models, leptin was associated with increases in both depression and fatigue. Leptin is known to implicated in linking obesity to depression^47^ and it is worth noting that BMI was elevated in our participant group (supplementary table 1).

### Altered brain connectivity in humans with PsD

We found that connectivity between the left thalamus and right nucleus accumbens, correlated positively with the PROMIS depression score in humans with PsD, as did connectivity between striatal and prefrontal areas. We also observed a positive correlation between fatigue/motivation and increased functional connectivity between right nucleus accumbens and right middle frontal gyrus, although this relationship was mostly driven by the association of fatigue/motivation and the depression score. Additionally, we found a non-significant but trending-positive correlation between thalamic amplitude of low-frequency fluctuation (ALFF - a proxy measure of spontaneous neuronal activity^21^) and depression score.

These data align with a recent human neuroimaging study that found higher connectivity between paraventricular thalamic nucleus and ventral striatum in individuals with anhedonia^22^. Associations between inflammatory markers and functional dysconnectivity in both discrete circuits and broad networks have been shown across numerous studies, as discussed in a recent review^48^. Another recent study^8^ used repeated imaging of individuals with depression and found that the volume of brain region forming the fronto-striatal salience network was almost doubled in cases of depression, and that this phenomenon was both stable across time and replicable across multiple cohorts. Interestingly, this network expansion was also seen prior to the onset of depression suggesting trait phenomenon. Considered alongside these other studies, our data suggest that neuroinflammation-driven depression seen in PsD engages the same neural circuits seen in other forms of anhedonia, and that different initial causes of anhedonia converge upon common neural pathways. The work from our group and others all indicate that midline thalamus, striatum and frontal regions should be key targets for therapeutic intervention.

### Neurobiological changes in reward circuitry

The mouse model of psoriasis-like disease afforded the opportunity to explore the neurophysiological response in detail and to characterise more fully the immunological response in the brain at a molecular and cellular level. The c-Fos expression in mouse within hours of Aldara application implicates midline thalamus as a key region in impaired reward/motivation behaviours. After 3 days of Aldara treatment, we found that the evoked thalamic AMPA receptor-mediated EPSC onto ventral MSNs was substantially weaker compared to control-treated mice, but with no change to the AMPA/NMDA ratio nor change in the frequency of amplitude of spontaneous EPSPs. The lack of change in AMPA/NMDA ratio would suggest the change in amplitude of the thalamic input is not a result of long-term depression or other forms of plasticity^49,50^.

Our interpretation of these data is that increased midline thalamic activity initially results in excessive excitatory neurotransmission. The precise mechanism through which Aldara treatment rapidly drives excitation in midline thalamus is unclear. However, data from a recent sheep study suggests TLR7 receptors may be present directly on thalamic neurons^51^, and we have previously shown that imiquimod directly enters the brain within hours of the first application^26^, both of which indicate a direct central effect of Aldara as opposed to secondary effects driven by peripheral inflammation. Inflammation decreases astrocytic clearance and buffering of glutamate by downregulation of glutamate transporters^52^, which may have exacerbated the consequences of increased thalamic excitability in striatum. Our observed downregulation of the astrocytic glutamate transporter *EAAT2/SLC1A2* transcript is consistent with this. Only 5% of striatal neurons are inhibitory interneurons^53^, far fewer than in cortex or hippocampus, so the lack of recurrent inhibition may make this region particularly vulnerable to excess glutamate. Indeed, Aldara treatment led to increased VGlut2-containing terminals in striatum but not cortex, with no change in VGlut1 immunolabelling, implying a specific alteration in the thalamostriatal pathway. We suggest that increased excitatory signalling leads to AMPA receptor internalisation in ventral MSNs. This is supported at the transcript level by reductions in Homer1, which is involved in AMPA receptor homeostatic scaling^54^, as well as downregulation of glutamate-gated ion channels^24^.

### Immune mechanisms underlying neuroinflammatory response

The main neuroinflammatory response in the rodent psoriasis-like model is likely to be mediated by astrocytes and microglia. We saw strong evidence of reactivity in these cells at both the gene and protein expression level and confirmed that both microglia and astrocytes were producing cytokines in our model. We found that the NLPR3 inflammasome was mediating at least some of the glial neuroinflammatory response. Activation of the NLPR3 inflammasome in microglia appears to be a key step for amplifying immune signals in the brain^30–32^ and NLPR3 has been posited as an important target in inflammation-driven depression^55–57^. We used a brain-penetrant NLRP3 inflammasome inhibitor^33^ in the Aldara model and found that this was sufficient to attenuate the production of cytokines by glia, suggesting that inhibiting NLRP3 may be able to act as an immune ‘circuit breaker’ and present a viable therapeutic target in immune-associated anhedonic depression.

While previous data indicate that microglia can proliferate in response to inflammation^58^, the apparent increase we seen post Aldara might be attributable to such proliferation, differentiation of ingressing monocytes, or perivascular macrophages. The absence of an increase, despite presence of reactivity, in the iCCR ^-/-^-mouse line could implicate the latter given their CCR2-dependent turnover. Immunohistochemistry revealed no morphological changes associated with astrocytes, using GFAP as a marker, yet a triple-positive marker approach to quantify reactive astrocytes with GFAP, S100β and ASCA2 via flow cytometry markers found these astrocytes significantly increased production of the proinflammatory cytokines IL-6 and IFN-γ. Given that the triple-positive subset of astrocytes that produced the proinflammatory cytokines constituted roughly a third of all S100β^+^ and ACSA^+^ astrocytes, we conclude that astrocytes had entered a reactive state despite GFAP immunohistochemistry showing no morphological change. Reactive microglia can induce reactivity in astrocytes via secretion of IL1α, TNFα, and C1q^29^, and subsets of reactive astrocytes can be defined by interferon-responsive genes such as *Igtp*, *Mx1* and *Ifit3*^59^. We found these genes to be significantly upregulated in treated *vs* control brains (see supplementary material).

While resident glial cells are likely to be a key mediator of neuroinflammation in our model, it is possible that infiltrating peripheral immune cells also contribute to the behavioural and circuit deficits either by acting as additional sources of cytokines in the brain or providing the initial inflammatory drive that is then amplified in the brain by glial cells acting via NLRP3-mediated pathways.. Recent studies have linked recruitment of peripheral monocytes into the brain with deficits in behaviour associated with inflammation^60^, with one study attributing cognitive impairments and neurite remodelling solely to circulating CX3CR1^+^ monocyte-derived TNF ^12^. We also detected CD4^+^ and CD8^+^ T cells throughout the brain parenchyma of Aldara-treated mice. Using different strategies to attenuate infiltration of the brain by either T cells or monocytes revealed that deficits in motivation cannot be attributed solely to either of these infiltrating immune cell types.

### Other considerations

It is worth noting that a recent study used the Aldara model and concluded that the model caused anxiety-like behaviours via IL17 receptor-expressing neurons in the amygdala, despite inhibition of these neurons not ameliorating the anxiety phenotype^61^. Of note, midline thalamus, particularly paraventricular thalamic nucleus, sends projections to the amygdala^62^, which in turn sends projections to ventral striatum^63^. The study by Lee *et al.*^61^ treated the Aldara model as a straightforward model of psoriasis but we previously found that the key component of Aldara, imiquimod, directly enters the brain to drive neuroinflammation via activation of TLR7. However, imiquimod (especially when formulated with isostearic acid in Aldara cream) can also cause a peripheral inflammatory response by driving a psoriasis-like skin inflammation^64^. As such, the neuroinflammatory response has two potential drivers: the direct effects of imiquimod (IMQ) on the brain and the effects of peripheral inflammation via neural and humoral routes, and these two may act synergistically^64^. It is not possible in this study to separate these two different inflammatory pathways.

The study by Lee and colleagues^61^ did not consider the brain penetrance of imiquimod and subsequent direct activation of resident glial cells, so it is likely that the mechanisms we revealed in the present study would, at least in part, explain their findings. Despite imiqiumod directly entering the brain, we do not believe that this is a significant confounding factor for interpreting our results, given that the thalamostriatal changes we have described in humans, as a consequence of inflammatory-mediated immune disease (psoriatic disease), indicate a relationship between systemic inflammation and neurobiological change. Additionally, the Aldara model is not chronic in contrast to the chronic inflammatory condition of PsD in the human data. It should be recognised that the inflammatory response in PsD is not consistently high and will be affected by the natural history of the disease and by treatment. However, our data could indicate consistency of thalamostriatal disruption irrespective of the duration of inflammatory exposure.

### Conclusions

We found that anhedonia/motivation behavioural changes occurring in the context of neuroinflammation are associated with perturbed thalamostriatial signalling in both humans and mice. This is indicative of species consistency in circuitry and relationship to behaviour. This is further supported by recent data describing primate murine homology in striatal inhibitory neurons.

These findings provide support for the validity of mouse models in studies of inflammation-induced neurobiological change and permit mechanistic studies impossible in humans. This circuit, and the immune mechanisms revealed in our study, present attractive targets for developing novel therapies, at least in the subset of depressed individuals showing heightened inflammation.

## Acknowledgements

We gratefully acknowledge the support of the Inger and George M Simpson Biological Psychiatry Scholarships. This work was supported by PhD studentships from the Saudi Arabia Ministry of Science (HAF) and the Government of Turkey (YK). The authors gratefully acknowledge the Glasgow Imaging Facility for their support and assurance in this work. The study was funded by Versus Arthritis (grant ref: 21964). The authors gratefully acknowledge the Flow Core Facility of the University of Glasgow for their support & assistance in this work. For the purpose of open access, the authors have applied a Creative Commons Attribution (CC-BY) licence to any Author Accepted Manuscript version arising from this submission.

## Material and Methods

### Human blood marker analysis

Sera samples obtained from 45 participants at baseline visits were processed and stored following a standardised protocol. Samples were thawed on wet ice or at 2-8 °C before use. To quantify the levels of circulating mediators we used an electrochemiluminescence multiplex assay (U-PLEX Metabolic Group 1 (hu) and S-PLEX Human IL-17A Kit, MSD Mesoscales Discovery, USA), which combines the classical sandwich immunoassay with electrochemiluminescence high-sensitivity detection techniques. To overcome cross-reactivity between proteins, analytes combinations in U-plex plates were carefully planned and optimised following producer’s recommendations. The signals generated by the standards are used to generate curve that fits a 4-parameter logistic model then used to determine the concentrations of circulating mediators.

For the blood marker analysis, we excluded markers where we did not have observations for all participants in our study. As the concentration of markers varied by orders of magnitude, we normalised all levels to the Z score. After normalising, several markers failed the Shapiro-Wilkes normality test so Spearman correlation coefficients were used. We used the FDR to correct correlation plots for multiple comparisons. All analyses and visualisations were carried out using R statistic software (v4.6.0; R Core Team 2026) in R Studio 2026 (Posit Team, Posit Software, PBC, Boston, MA) with *tidyverse* ^65^*, dyplr* and *ggplot2* ^66^ packages. Elastic net regression was run using the *glmnet* package ^67^ in R. Correlation plots were generated using the *corrplot* package (Wei & Simko, 2024, v0.95; https://github.com/taiyun/corrplot). All data and code for these analyses can be downloaded from https://github.com/MickCraigLab/inflammation-thalamostriatal.

### Human neuroimaging

In an observational study with patients living with PsD, we used 3T MRI to characterize the thalamo-striatal functional connectivity association with depression and motivational impact of fatigue. The clinical cohort (n=46) had a mean age of 49 years (SD 11.2), was 52% female, and demonstrated high to moderate disease activity (mean Disease Activity in Psoriatic Arthritis [DAPSA] 40.8, SD 19) with an average disease duration of 6.4 years (SD 5.8). Depressive symptoms were assessed using the standardised PROMIS Depression short form: 68% met criteria for depressive symptoms, including 13% classified as severe, 15% moderate, and 30% mild. Fatigue was also systematically evaluated using both numeric rating and the standardized PROMIS Fatigue short form16. The mean Fatigue NRS was 6.5 (SD 2.6), and the average of PROMIS Fatigue score was 51.3 (SD 16.6), indicating a moderate burden of fatigue. To further characterize its functional impact, we included subcomponents of the PROMIS Fatigue measure, focusing on motivational aspects. The PROMIS Fatigue motivational impact subscore averaged 13.4 (SD 4.2), while responses to individual items specifically addressing motivational fatigue, namely M3 and M4, yielded mean scores of 3.3 (SD 1.0) and 3.7 (SD 1.0), respectively (see supplementary table 1).

We defined bilateral nucleus accumbens and thalamus based on the Harvard-Oxford Atlas. Functional connectivity (FC) was calculated as Fisher z transformed Pearson correlation coefficients between the time series of BOLD signal changes between bilateral thalamus and bilateral nucleus accumbens after preprocessing using the default CONN toolbox pipeline^68^. Linear regression was used to demonstrate an association between thalamocortical connectivity and depression in PsD, after controlling for age and sex.

We computed brain maps of amplitude of low-frequency fluctuations (ALFF) in the CONN toolbox to determine the baseline activity in thalamostriatal regions in relation to depression. These maps represent a measure of BOLD signal power within the frequency band (0.008–0.09 Hz) and are defined as the root mean square of BOLD signal at each individual voxel^21^. ALFF values were normalised for each patient, thus the distribution of ALFF values across all voxels is normal. This allows the normalised ALFF values to show the relative amplitude compared to other areas: positive values indicating more active compared to average activity, negative values indicating less active. These ALFF values were then used in linear regressions with depression scores after controlling for age and sex to determine any association between thalamostriatal activity and depression in PsD. Ethical approval was granted by the West of Scotland Research Ethics Committee (ref 19/WS/0033), and all participants provided written informed consent in accordance with the Declaration of Helsinki.

### Mice

Female C57BL/6J mice were obtained from Charles River Laboratories (UK) and housed in individual ventilated cages. Animals underwent acclimatisation for one week prior to experimental use at 8 to 10 weeks of age (group numbers are described per experiment below). Animals were maintained in a 12 h light/dark cycle in controlled temperature and humidity with *ad libitum* access to food and water. All experiments were carried out in accordance with the UK Animal (Scientific Procedures) Act 1986 and were subject to local ethical approval by the University of Glasgow Animal Welfare and Ethical Review Board. The iCCR^-/-^mice, with genes deleted for four inflammatory chemokine receptors *CCR1, CCR2, CCR3 and CCR5 (iCCRs),* were obtained from the Chemokine Research Group within the University. These animal strains were generated and maintained on a C57BL/6 background and bred in a specific pathogen-free environment.

### Aldara model

C57BL/6, and iCCR^-/-^female mice (8 to 22 weeks old, starting weight range: 16.3 – 29.5 g) were each treated with a ∼62.5 mg dose of Aldara™ (Meda AB, UK) cream containing 5% Imiquimod - IMQ or an equivalent volume of control cream (Boots Aqueous Cream B.P.; Boots Pharmacy, UK) by topical application to shaved dorsal skin near the base of the tail (∼3 mm^2^ area). Treatment was applied for 3 consecutive days, with tissue collected 24h after the final application. Soft diet was provided to both control and experimental groups and mice monitored daily for weight loss. Female mice only were used in this study due to male mice showing differences in inflammatory response to topical application of inflammatory agents^34^.

### Experimental design

Mice were randomly allocated to treatment groups. A power analysis was performed using G*Power ^69^ software (v3.0.10 G*Power, RRID:SCR_013726) to determine minimum group size; n

= 4/5 (immunostaining), n = 4 (flow) and n = 8 (behaviour) were selected for each treatment group. The effect size for output was based on previous or pilot experiments, with calculations made on a basis of 80% power, and a significance criterion of 0.05. All tissue was coded prior to imaging and analysis to prevent researcher bias.

### Behavioural assessment

#### Animals

Following acclimatisation, they were singly housed in rat cages (previously washed and sterilised to eliminate any other species’ scents), equipped with a running wheel and two water bottles, one of which contained 0.1% (w/v) sucrose solution, available *ad libitum*, to which the mice were acclimatised. Behavioural testing was conducted throughout (nest-building, sucrose preference) or on day 4, 24 hours after the last application of cream. Elevated plus maze was conducted in the morning and mice were returned to their cage room for 2 – 3 hours before performing open field test in the afternoon. Before each test, mice were allowed to acclimatise to the behaviour room for 30 mins prior to the start of testing.

#### Sucrose preference

Mice were allowed to establish their pattern of consumption of water and sucrose over the course of several days, prior to the treatment with topical creams. Both water bottles were weighed in the mornings and evenings and the change in weight was compared as a ratio of sucrose preference.

#### Nest building

We used the Deacon nest-building protocol^25^. Pressed unbleached cotton squares of 3 g (± 0.05 g) each were given every morning for nest building and collected the following morning. The images of the nests were taken for rating out of 5 (where 5 points were given to a fully shredded cotton material built up into a high walled nest, covering the mouse; 1 point reflects an untouched pressed cotton material) in the evenings as well as following mornings. The non-shredded material was weighed as another measure of nest building.

#### Elevated plus maze

To assess anxiety, mice were placed into cross shaped Perspex arena (110 cm x 110 cm, 65 cm from ground) for one trial of 5 mins only and recorded using ANY-maze video-tracking software (ANY-maze, Ireland). ANY-maze software was also used to analyse the following: distance travelled, average and maximal speed in different parts of the arena (closed arms *vs* open arms) as well as numbers of entries and time spent in open arms (OA) versus closed arms (CA) of the arena.

#### Open field test

To assess locomotor activity and as a secondary measure of anxiety in a novel arena environment, mice were placed into individual Perspex boxes (40 x 40 x 45 cm) for 1 h and recorded using ANY-maze video-tracking software. First 15 mins were analysed using ANY-maze software, and measures recorded included distance travelled, average and maximal speed in different parts of the arena as well as numbers of entry and time spent in centre and perimeter of the arena.

#### Rotarod

We measured locomotor activity using an ITC Rotarod (IITC Life Sciences Inc, USA). Mice were habituated to the room for 30 min prior to testing, and acclimatised to the accelerating rotarod (0 – 60 rpm) for 3 trial runs the day before recording. A baseline performance was recorded immediately before the first topical cream application. Mice were placed on the beam in separate lanes of the rotarod apparatus. Latency to fall from the rod was recorded for up to 300 s each trial for three trials per day on 3 consecutive days. The latency to fall compared to the baseline was reported per mouse and compared between groups.

### Stereotaxic surgeries

Several midline thalamic nuclei target ventral striatum including but not limited to paraventricular (PVT), centromedian (CM) and parataenial (PT) thalamic nuclei^70^. To broadly target midline thalamus for optogenetic and electrophysiological experiments, we carried out stereotaxic injections to deliver 300 nl of AAV8-hSyn-Chronos-GFP (UNC Viral Vector Core, USA; contributed by Ed Boyden; titre 3.1 × 10^13^ viral particles / ml) to the border of CM and PVT. We used the following stereotaxic coordinates: R/C −0.5 mm and M/L 0 mm from Bregma, at a depth of 3.3 mm from the pial surface. Briefly, mice were anaesthetised with 5% Isoflurane, were placed on a heated pad for the duration of the surgery and maintained at 1.5–2.5% isoflurane (with a flow rate of ∼2 l min^−1^ O_2_). Mice were then subcutaneously dosed with 0.1 mg/kg of buprenorphine (buprenorphine hydrochloride, Henry Schein) and 5 mg/kg of carprofen (Rimadyl, Henry Schein) at the start of surgery, and additional doses of carprofen were provided subcutaneously subsequent days, as required. Anaesthetised mice were placed in a Kopf 940 Small Animal stereotaxic frame (David Kopf Instruments, CA, USA). An incision was made down the midline, and a craniotomy was performed to allow injection of virus (300 nl) using a Neuros Syringe (Hamilton Company, NV, USA) injected at a rate of 100 nl/min using a UMP3 UltraMicroPump (World Precision Instruments, UK). Mice were recovered for 3 weeks to allow sufficient time for opsin expression to occur before undergoing Aldara treatment as described above.

### Electrophysiology experiments

#### Drugs and chemicals

CGP55845, DNQX, gabazine, DL-AP5 and picrotoxin were purchased from either Tocris Bioscience (UK) or HelloBio (UK), and all other chemicals were purchased from Sigma-Aldrich (UK) unless otherwise stated.

#### Brain slice preparation

Mice were deeply anesthetised with 5% isoflurane and the brain was rapidly dissected out in room temperature NMDG cutting solution, containing (in mM): 135 NMDG, 20 choline bicarbonate, 10 glucose, 1.5 MgCl_2_, 1.2 KH_2_PO_4_, 1 KCl, 0.5 CaCl_2_, saturated with 95% O_2_ and 5% CO_2_ (pH 7.3-7.4). Brains were hemisected and coronal slices (400 um) were cut using a VT-1200S vibratome (Leica Microsystems, Germany) and the slices were transferred into a chamber containing recording aCSF, composed of (in mM): 130 NaCl, 24 NaHCO_3_, 3.5 KCl, 1.25 NaH_2_PO_4_, 2.5 CaCl_2_, 1.5 MgCl_2_, and 10 glucose, saturated with 95% O_2_ and 5% CO_2_ (pH 7.4). Slices were placed in a water bath at 37 °C for 30 minutes and slowly cooled to room temperature for further 30 minutes, at which they were maintained until recording.

#### Patch-clamp recording

For *ex vivo* patch clamp recordings, slices were attached onto 0.1% poly-L-lysine (Sigma Aldrich, P8920) coated glass slides and placed into the upright microscope and visualized using infrared oblique contrast microscopy (Scientifica SliceScope, Scientifica, UK). A CoolLED pE-4000 system (CoolLED, UK) was used to confirm presence of thalamic axons in slices, and to provide optogenetic stimulation. The slices were submerged in recording aCSF, warmed to 32-34 °C, and the rate of perfusion was kept at 5 ml/min. The recording electrodes resistance was typically 4 – 6 MΩ and were pulled from borosilicate glass (World Precision Instruments, UK) using a Sutter P-1000 horizontal puller (Sutter Instruments, CA, USA). The intracellular solution used had the following composition (in mM): 135 Cs-methanesulphonate, 8 NaCl, 10 HEPES, 0.5 EGTA, 4 MgATP, 0.3 Na-GTP, 5 QX314, plus 2 mg/ml biocytin (VWR International, UK), at pH 7.25 adjusted with CsOH to 285 mOsm. Whole-cell recordings in voltage clamp mode were made from ventral striatum, targeting medium spiny neurons (MSNs) based on morphology. Gabazine (10 µM) and CGP55845 (1 µM) were present in the bath throughout recordings to block GABA_A_ and GABA_B_-receptor-mediated currents, respectively.

Upon formation of a whole-cell patch, spontaneous activity was recorded for 1 to 2 minutes to sample spontaneous EPSC activity, at a holding potential of −70 mV. A single light pulse was of 470 nm was then applied once every 10 seconds to optogenetically-excite midline thalamic inputs onto ventral MSNs. After a suitable baseline had been established (typically >20 sweeps), 10 µM of AMPA receptor antagonist DNQX was added to the bath to inhibit AMPA-mediated synaptic currents. The holding potential was then switched to +50 mV to allow recording of NMDA receptor-mediated synaptic currents. To confirm the presence of the NMDA current, D-AP5 (100 µM) was added at the end of the recording in a subset of recordings. Whole-cell patch-clamp recordings were made using a Multiclamp 700B amplifier (Molecular Devices, CA, USA). Signals were low-pass filtered at 8 kHz and digitized at 20 kHz using a Digidata 1440A and pClamp 10.2 (Molecular Devices, CA, USA). Recordings were not corrected for the liquid junction potential.

#### Data Analysis

Electrophysiology data were analysed in ClampFit 11 (Molecular Devices, CA, USA). To measure spontaneous EPSCs, traces were first band-pass filtered in using a high-pass 8 pole Bessel band pass filter at 2Hz and a low-pass Gaussian filter at 1000Hz, and sEPSCs were detected using template matching. For evoked EPSC analysis, data were first baselined to zero and an average was taken of at least 20 traces for both AMPA-and NMDA-receptor-mediated components. The AMPA receptor-mediated EPSC was determined as the maximal EPSC peak at −70mV and NMDA receptor-mediated EPSC as the highest peak at +50 mV. For example trace figure preparation, recordings were imported into Igor Pro (Wavemetrics, OR) using Neuromatic (Thinkrandom, UK). Raw sEPSC traces were band-pass filtered in Igor Pro between 0.1 and 1000 Hz to improve clarity. GraphPad Prism (Graphpad, CA) was used for statistical analysis. Data were tested for normality using the D’Agostino and Pearson test and subsequently analysed by parametric or nonparametric tests as appropriate. Unless otherwise stated, all values are presented as mean ± SEM.

#### Post hoc morphological recoveries

The tissue slices were post-fixed in 4% PFA solution for an hour after recordings and then transferred to PBS (Melford, UK). Slices were washed thrice in 0.1 M PBS solution, followed by three further washes in PBS with 0.5% Triton X (Sigma, UK). Two blocking steps were used: 100mM of Glycine (Sigma, UK) incubation for 20 mins and then 1-hour long incubation with blocking buffer with 10% goat serum (VectorLabs, UK) at room temperature. Primary antibody against GFP (Abcam, UK) at 1:5000 dilution was used overnight at 4°C; then secondary conjugated anti-chicken antibody (ab150173, Abcam UK) and streptavidin (VectorLabs, UK) were used at 1:250 and 1:500 dilutions respectively were used in a carrier solution containing 10% goat serum and PBS for a two-hour incubation at RT. Finally, the slices were washed thrice in PBS and transferred to 30% sucrose (Thermofisher, UK) until cryoprotected. Then slices were re-sectioned at 100-micron thickness using SM2010R freezing microtome (Leica, UK)) at −20°C and mounted onto glass-slides and HardSet mounting medium with DAPI was used (VectorLabs, UK).

### Immunohistochemistry

Mice were given a terminal overdose of the anaesthetic sodium pentobarbitone (Euthatal, 10 μl/g of 200 mg/ml solution) and, once deeply anaesthetised, were killed by transcardial perfusion with PBS followed by 4% paraformaldehyde (Sigma-Aldrich, UK). Following perfusion-fixation, whole brains (n=4 / treatment) were dissected into 4% PFA overnight at 4°C, then cryoprotected in 30% sucrose at 4°C. Coronal sections were cut at 40μm on a SM2010R freezing microtome (Leica, UK) and collected into in-house solution of antifreeze (containing 30% ethylene glycol, 25% glycerol and 25% 0.2M PB in ddH_2_O) and stored at −20°C until required. Free-floating sections were washed in PBS then treated with 100% EtOH at −20°C for 10 min, or an antigen retrieval method (overnight at 60°C in 10 mM sodium citrate buffer (pH 6.0)). Tissue was thoroughly washed in PBS, then blocked in PBS containing 10% normal goat serum (NGS) + 0.3% Triton X for 1h at 4°C. Primary antibodies were prepared in PBS containing 3% NGS and applied overnight (or 72 hours in the case of IBA-1, GFAP, VGLUT1 and VGLUT2) at 4°C. Primary antibodies included a combination of the following: rat anti-CD4 antibody (Biolegend, UK, cat# 100505, 1:500), guinea pig anti-IBA1 antibody (Synaptic Systems, Germany, Cat# 234 308, 1:500), rabbit anti-GFAP antibody (Novus Biologicals, UK, Cat# NB300-141, 1:500), rabbit anti-c-fos (Cell Signalling Technologies, UK, cat # 2250S, 1:500), rabbit anti-VGLUT1 (Synaptic systems, Germany, Cat# 135-302, 1:500), guinea-pig anti-VGLUT2 (Synaptic systems, Germany, Cat# 135-418, 1:250). Sections were washed 3x 10mins in PBS, before the appropriate combination of secondary antibodies were applied for 2h at RT. Secondary antibodies were as follows: isotype-specific Alexa Fluor 488- and Alexa Fluor 546-conjugated goat anti-rat IgG antibodies (Thermo Fisher Scientific Cat# A-21434, RRID:AB_2535855; 1/500), Alexa Fluor 488 goat anti-guinea pig (Abcam, UK, Cat# ab150185, 1:500), Alexa Fluor 594 goat anti-rabbit (Invitrogen, USA, Cat# A11012, 1:500). 3 x 10min PBS washes were followed by mounting in VECTASHIELD® HardSet™ mounting medium with DAPI (Vector Labs, UK, Cat# H-1500-10).

### Anatomical tracing of the thalamostriatal projections

Mice that had undergone stereotaxic injections of viral vectors to transduce Chronos-GFP (details below) were killed by perfusion-fixation as described above. Coronal sections were sliced at 50 µm. The slices around the injection coordinates as well as positive control areas (such as prefrontal cortices) and experimental striatal slices were selected and underwent the immunofluorescent staining protocol described above. To enhance visualisation of thalamic axons, sections were incubated with primary antibody against GFP (chicken anti-GFP; Abcam, UK, cat # ab64905, 1:50000) followed by incubation with Alex488-conjugated goat anti-chicken secondary antibody (Abcam, UK, cat # ab150173,1:250). Once stained, the slices were mounted using the HardSet mounting medium containing DAPI (VectorLabs, UK), and images were taken on SLM710 confocal microscope (Zeiss).

### Image acquisition and analysis

For immune cell infiltration studies, tile scans of entire coronal brain sections were captured using a widefield inverted Leica DMi8 microscope with a 10x objective. FIJI (ImageJ) software^71^ was used to analyse the images as follows: the DAPI channel was used to outline each selected brain region with reference to the Allen brain atlas. The number of CD4^+^ cells within this measured area were counted per mouse, the mean plotted and compared per treatment for each region. Criteria for cell inclusion included CD4^+^ signal, a DAPI nucleus and a diameter of 8-12 µm. Illustrative images were captured using a Zeiss LSM 880 Axio Imager 2 laser-scanning confocal microscope (Zeiss Microscopy, UK) processed using Zen Zeiss Microscopy Software.

For c-fos quantification in the thalamus and VGLUT analysis in the ventral striatum and barrel cortex, confocal z-stacks were captured using a Zeiss LSM 880 Axio Imager 2 laser-scanning confocal microscope equipped with an LD C-Apochromat 20x/1.1 W Korr M27 objective at 0.8 µm optical slice and Plan-Apochromat 63x/1.4 Oil DIC M27 objective at 0.4 µm optical slice, respectively. The same exposure levels were used to capture images from all sections. Quantification of the number of fos-positive cells per ROI was performed in FIJI from maximum intensity z-projections. Quantification of VGLUT positive terminal number and size was performed in FIJI from three 50×50 µm ROIs taken from single z-slices of three sections per mouse.

For analysing the fluorescence signal intensities and coverage areas of IBA-1 and GFAP, confocal z-stack images of IBA-1 and GFAP immunofluorescence were acquired using a Zeiss LSM 880 Axio Imager 2 laser-scanning confocal microscope equipped with an LD C-Apochromat 40x/1.1 W Korr M27 objective (Zeiss, 421867-9970-000) and Immersol W 2010 immersion oil (Zeiss, 444969-0000-000). Each image stack consisted of 16 optical sections spaced 2 µm apart (total depth 30 µm) with a resolution of 2048 x 2048 pixels (354.25 µm x 354.25 µm).

To quantify IBA-1 and GFAP signal intensity and coverage area, the images were first processed the images to subtract the background signals using a 35-pixel rolling ball radius applied to each z-stack in Fiji. Maximum intensity z-projections were then generated, and positive signals isolated using pre-trained pixel classifiers for IBA-1 and GFAP within the LabKit Fiji plugin^72^. The resulting binary masks were used to calculate the percentage area covered by positive signal and to redirect measurements to the original image to measure the mean grey value within the positive signal regions.

### Whole brain transcriptome analysis

We carried out a secondary analysis of whole-brain RNAseq transcriptomic data reported in detail elsewhere^24^. In brief, differential expression analysis was carried out DESeq2^73^ using pairwise models with no additional covariates. The normalised data were explored using the Searchlight analysis platform^74^. The significance threshold for differential genes was adjusted p < 0.05 and absolute log2fold > 0.5. All other parameters were left to default.

### Flow cytometry

Mice were given a terminal overdose of the anaesthetic sodium pentobarbitone (Euthatal, 10 μl/g of 200 mg/ml solution) and, once deeply anaesthetised, were killed by transcardial perfusion with 20 ml of ice-cold PBS. Whole brains were dissociated using the Adult Brain Dissociation Kit for mouse (Miltenyi Biotech: Cat. No.:130-107-677, UK) in line with manufacturer’s instructions. Briefly, whole brains were cut into 8-10 pieces and placed into a C-tube containing the enzyme mix. Samples were dissociated in the gentleMACSTM Octo Dissociator with heaters using program 37C_ABDK_01 (Miltenyi, UK). Subsequently, samples were filtered through a 70 μm strainer using 10 ml of D-PBS and the back of a 5 ml syringe plunger before being spun at 300 g for 10 min, 4°C. The supernatant was discarded, and cells were re-suspended in 3.1 ml D-PBS and 900 μl Debris Removal Solution. Another 4 ml of D-PBS was overlain gently, and samples were spun at 3,000 g for 10 min, 4°C, with half (5) acceleration and half (5) brake. The top two phases, containing debris and myelin, were removed and discarded. D-PBS was added, and cells were centrifuged at 1,000 g for 10 min, 4°C, and supernatant discarded. Erythrocytes were removed by incubating with 1X Red Blood Cell Removal Solution (provided in the kit) diluted in water for 10min at 4°C before adding cold PB Buffer. Samples were spun again at 300 g for 10 min, 4°C and the pellet was dissolved in 1 ml of PBS. Trypan blue and hemacytometer were used for cell counting.

Brain cells were stained in 96-well plate, following protocol with FACS (2% FBS and 0.4% of 50 mM EDTA in 1X PBS) buffer wash and centrifuge spin (400g for 5min at 4°C) in between each step (all incubations were performed at 4°C unless stated otherwise). Live/dead staining (eBioscience™ eFluor™ 506 Catalog number: 65-0866-14, USA) was done in PBS for 20 mins. Fc block (Miltenyi, 130-059-901, UK) step was done for 20 mins and conjugated extracellular antibody (see supplementary table 2) staining was done in FACS buffer for 20 mins. For intracellular staining from T cells, single cells, post extracellular staining, were stimulated with cell stimulation cocktail with protein transport inhibitor (eBioscience™ Cell Stimulation Cocktail (plus protein transport inhibitors) (500X) cat. No. 00-4975-03, USA) for 4 hours at 37 C° in CO_2_ incubator. Cells were fixed (Fix buffer, BD Biosciences, USA cat no. 554655) and permeabilised with Perm buffer (BD Biosciences, USA cat. No. 554723) and intracellular staining (see supplementary table 2) was done in perm buffer. For cytokine staining from microglia, astrocytes, and monocytes, only Golgi blocker, Brefeldin, was used for 4 hours. All fluorescence minus one (FMO) staining controls received the full panel staining except for one antibody to determine true populations in the sample (see supplementary table 2) FMOs were generated using pooled sample cells. Compensation was performed with antibody stained UltraComp eBeads (Thermo Fischer Scientific, USA, cat No. 01-2222-41). Flow cytometry was performed at the Cellular analysis facility, MVLS shared research facilities (University of Glasgow) on the LSR Fortessa (BD Biosciences, USA). Data is acquired through BD FACSDiVaTM digital software. All analysis was performed using FlowJo software (v.10.10.0.) (BD Biosciences, USA). Graphs were generated using absolute cell number using the cell counts from haemocytometer and gating strategies percentages (see supplementary methods).

### T Cell Depletion experiment

Four cohorts of female C57BL/6J mice (n=4, each cohort) were used for T cell depletion experiment. 2 cohorts were injected with 400 µg of rat IgG (LTF-2) and another 2 cohorts with depleting 200 µg CD4 (GK1.5) and 200 µg CD8 (2.43) antibodies, respectively intra peritoneum (*ip*), three days before starting the control and Aldara cream treatment. Another set of these blocking antibodies were administered i.p. couple of hours before the 2nd day of cream treatment. Tissue collection was performed after the final (3^rd^ day) cream application, as described above.

### Immunoblotting

Whole brain homogenates were prepared using RIPA buffer with protease and phosphatase inhibitor. Samples were run on 4 - 12% gradient gel and transferred to nitrocellulose membrane. 5% milk in TBS-tween was used for 1hr blocking. Anti IL-1b (mouse, 1:1000) and NLRP3 (rabbit, 1:1000) were used with anti-mouse secondary antibody. Α-tubulin was used as loading control.

### NLRP3 inhibition

We used a brain penetrant NLRP3 inhibitor^33^, NT-0527 (NodThera, UK) at 100 mg/kg in a vehicle composed of 0.5% methyl cellulose (cat. # M7410; Sigma-Aldrich, UK) and 0.2% Tween 80 (cat. #P4780; Sigma-Aldrich, UK). Mice were dosed with drug (20 mg/ml) or vehicle via oral gavage, starting one day prior to Aldara treatment, and continued for all three days of the treatment, an hour before applying the cream. Tissues were harvested after final day (3^rd^ day) of cream application, as described above.

### Statistical analysis

Data were checked for normality using Shapiro-Wilk’s test and parametric tests were used throughout, unless otherwise specified; outliers were tested for using ROUT test. No animals were excluded from the data analysis. Comparisons were made using either unpaired two-tailed t-test (e.g. For total average speed in EPM maze for control and Aldara-treated animals) or one-way ANOVA with *post hoc* multiple comparisons with Tukey correction. Data are presented as mean ± SEM unless otherwise stated. Unless otherwise stated, the numbers of independent animals are described per experiment above and indicated in the figure legends. Statistical differences were determined using GraphPad Prism software (GraphPad Prism, RRID:SCR_002798).

