## Supplementary data for "Psoriasis-related neuroinflammation disrupts thalamostriatal signalling driving anhedonia in both humans and mice"

### Supplementary Figures

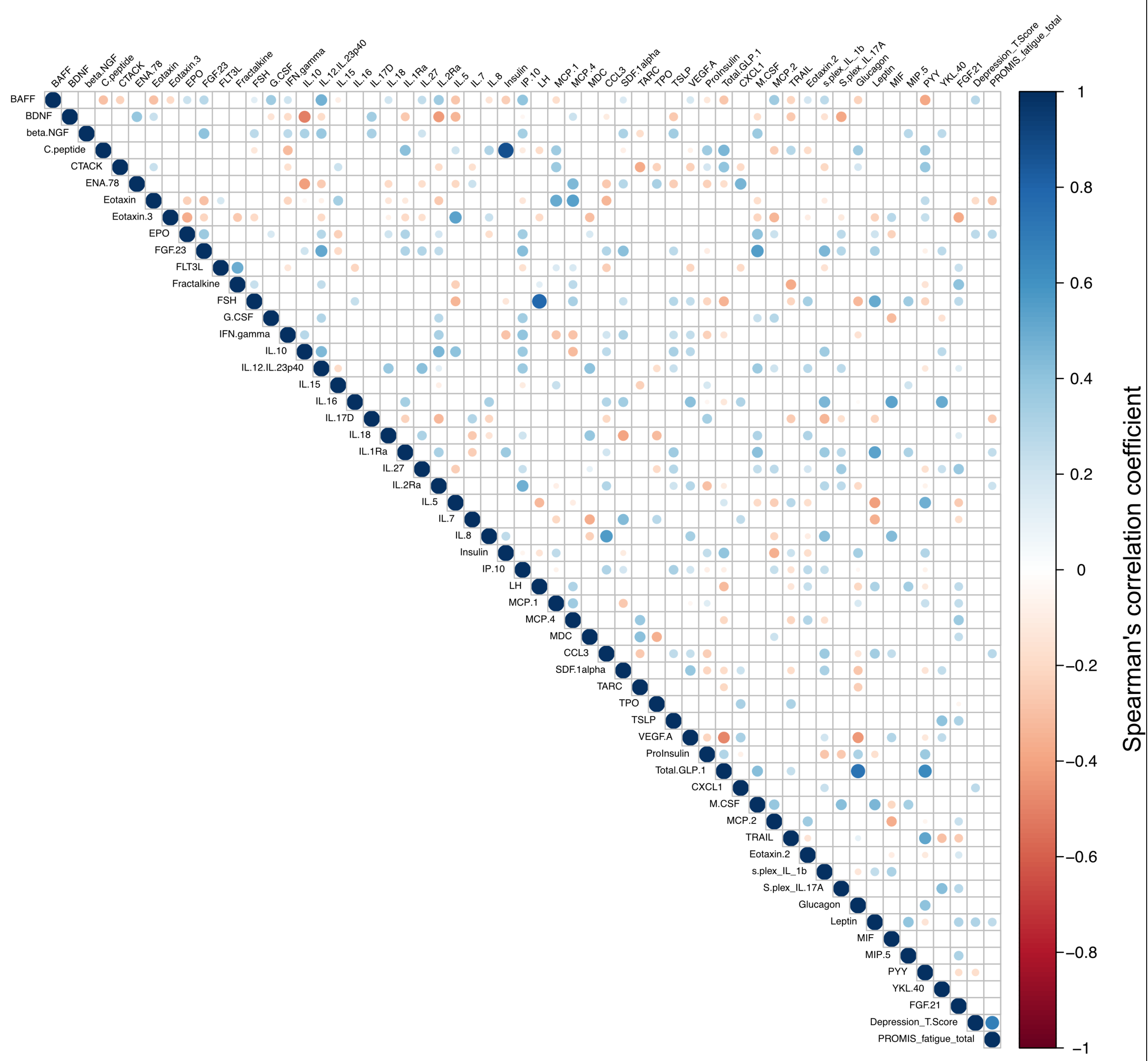


**Supplementary figure 1: Correlation plot of electroluminescence immunoassay markers with depression and fatigue scores.** Each dot represents the Spearman's correlation coefficient between each factor tested. We only included analytes for which we had data for all participants (n=45). Positive correlations shown in blue, and negative correlations in red, with size of point and intensity of colour both reflecting the magnitude of the correlation coefficient. Non-significant correlations ($p < 0.05$ after correcting for FDR) have been excluded from the plot.

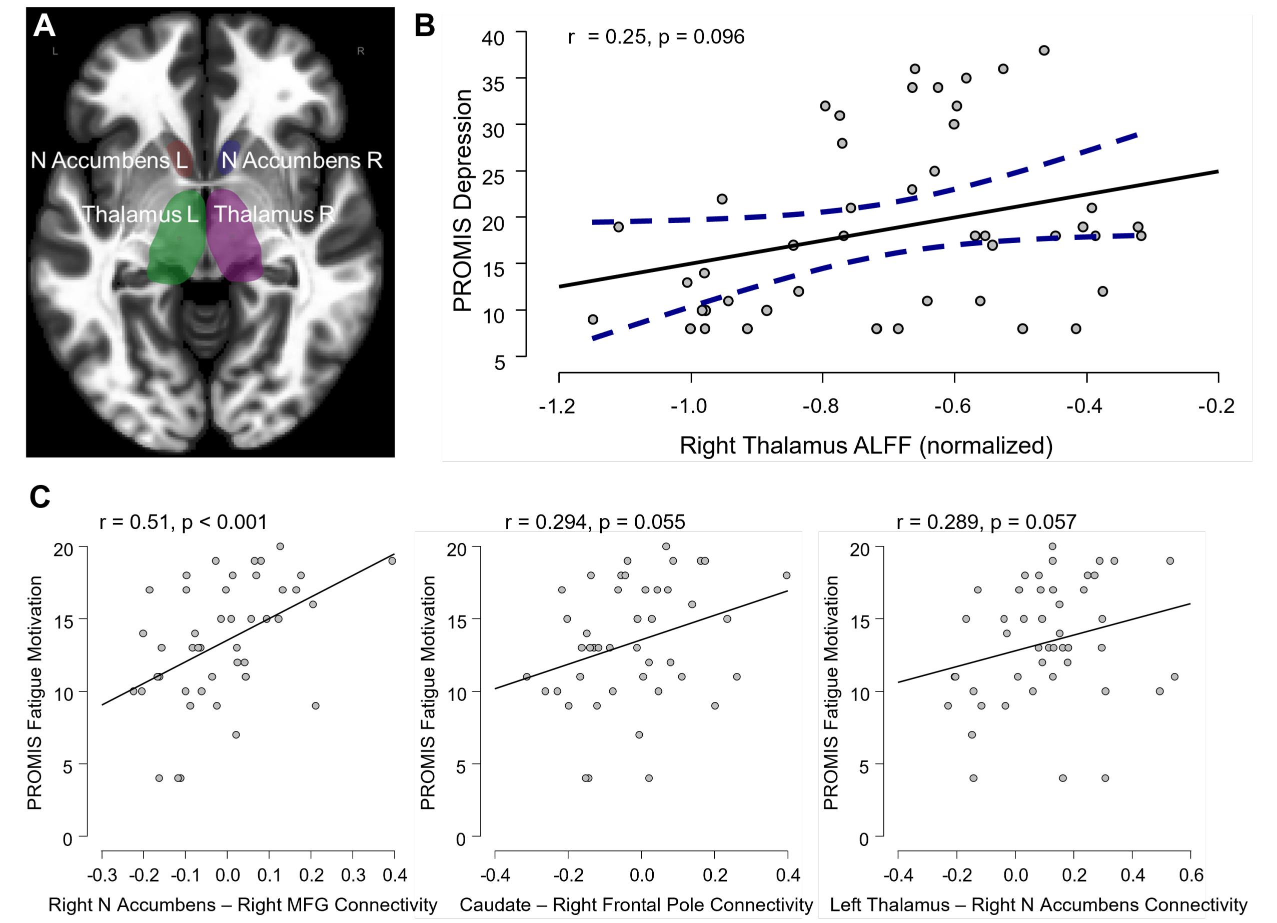


**Supplementary figure 2: Additional neuroimaging data. A**, an axial slice (x = 10, y = -18, z = -6) of an MNI template with the four regions-of-interest (ROIs): the bilateral nucleus accumbens and thalamus, based on the Harvard-Oxford Atlas. **B**, spontaneous functional activity of the right thalamus shows a trend towards significance of positively correlating with PROMIS Depression scores (r = 0.25, p = 0.096) using normalized amplitude of low-frequency fluctuations (ALFF) values. **C**, We saw a significant correlation for connectivity between right nucleus accumbens and right middle frontal gyrus (MFG) with the PROMIS fatigue score and borderline significance correlations with connectivity between caudate and frontal right pole, and left thalamus and right nucleus accumbens. However, when treating the PROMIS depression score as a factor in the model, the significance of the correlation disappeared for all of these regions, indicating that depression is the key driver of fatigue and brain connectivity. All analyses are controlled for age and sex.

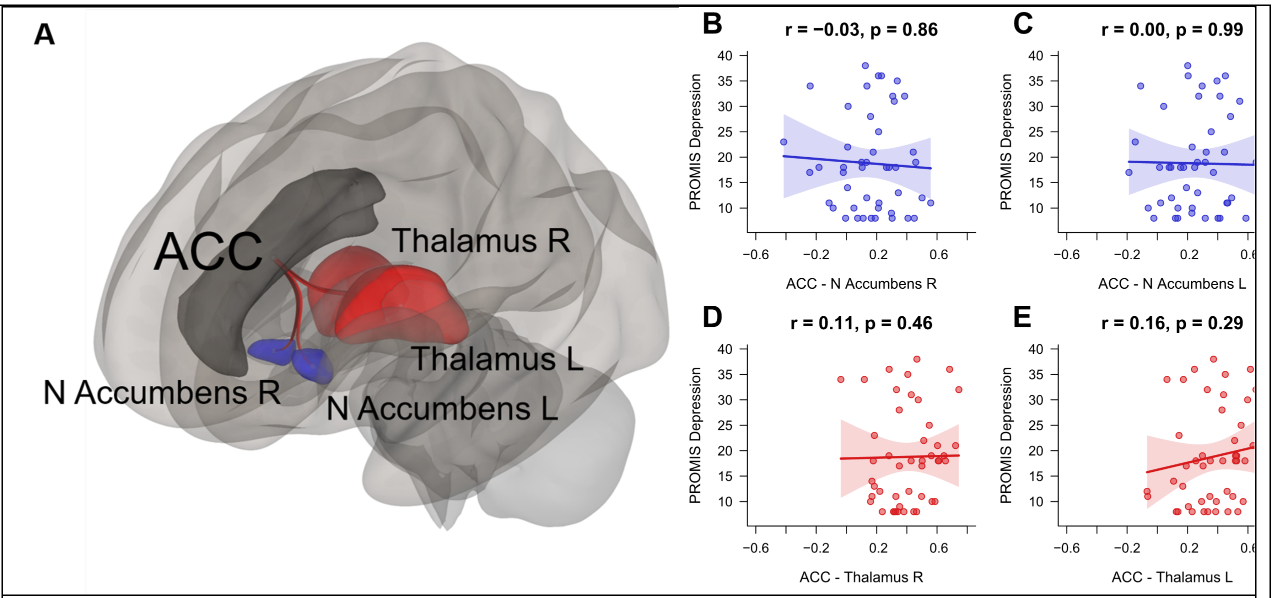


**Supplementary figure 3: functional connectivity of anterior cingulate cortex (ACC) does not correlate with depression. A**, schematic of connectivity tested. Functional connectivity between ACC and right (**B**) or left (**C**) nucleus accumbens did not correlate with the PROMIS depression score. Similarly, functional connectivity between ACC and right (**D**) or left (**E**) thalamus did not correlate with the PROMIS depression score.

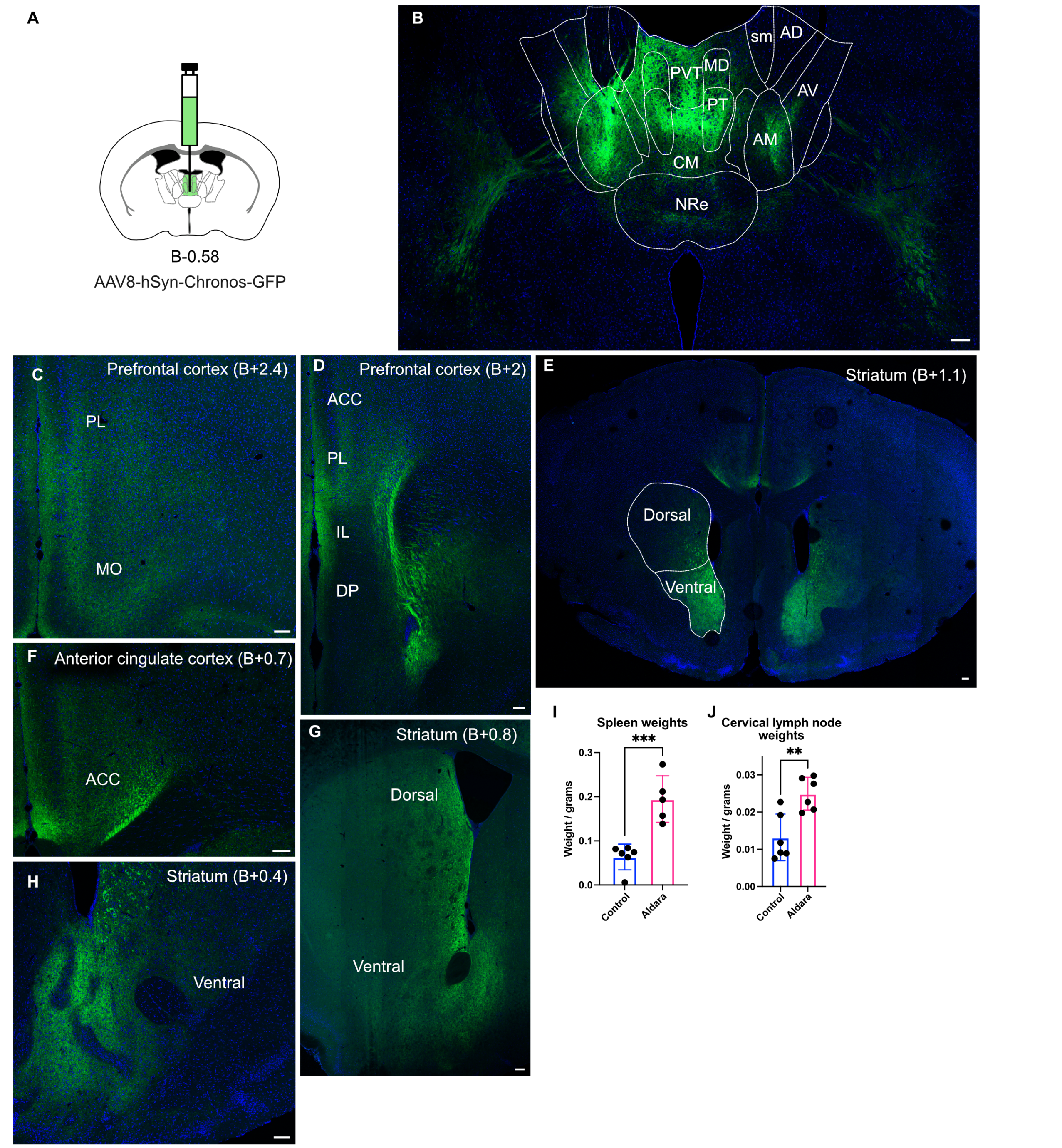


**Supplementary figure 4: Additional images confirming axonal projections from midline thalamus from optogenetics experiments. A**, schematic of target injection site. **B**, representative image of viral spread in the injection site. **C – D & F**, representative images of fibres present in the prefrontal cortices, including all of the subfields at different points on rostro-caudal plane (coordinates relative to Bregma). **E**, typical spread of fibres in the striatum and the delineation of dorsal vs ventral striatal areas. **G & H**, typical fibre distributions in striatum at different points on rostro-caudal plane. **I & J**, spleen and cervical lymph node weights, showing an increase in both weights in the Aldara-treated mouse groups in mice used for electrophysiology; statistical comparisons are made with student's *t* test; **, $p < 0.01$; ***, $p < 0.005$.

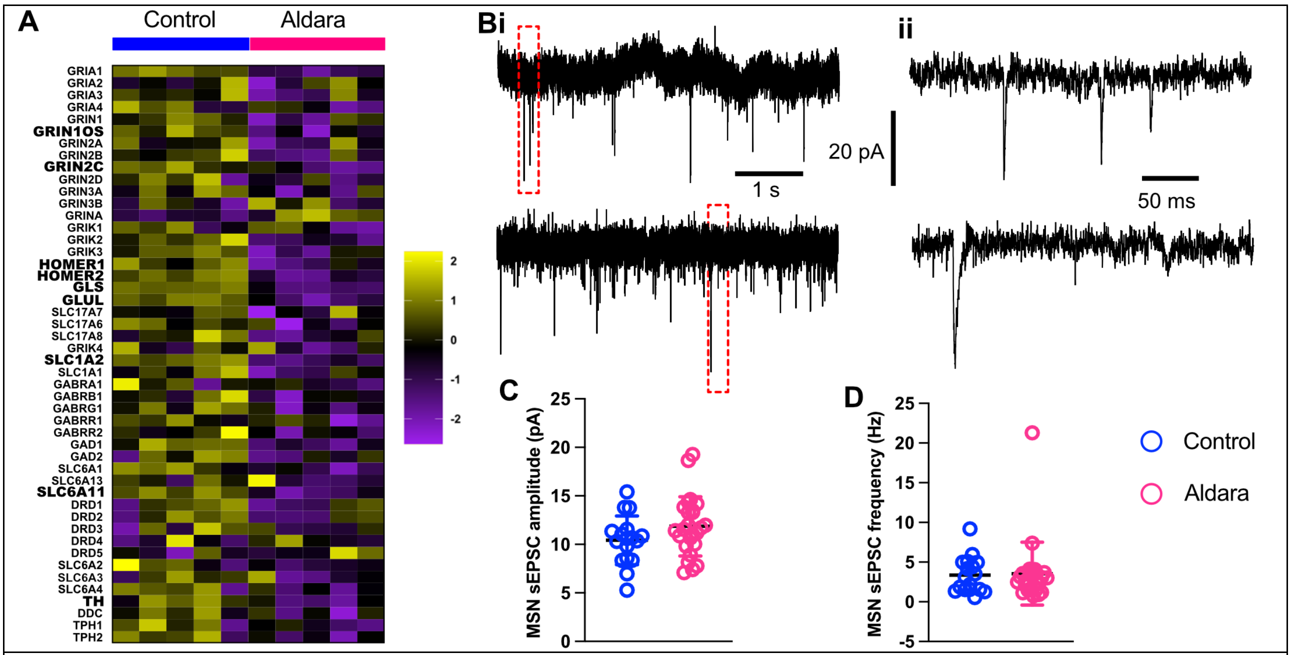


**Supplementary figure 5: Baseline neurotransmission after Aldara treatment. A**, heatmap showing bulk RNAseq analysis for a selection of neurotransmission-related genes for whole brain transcriptome for control-treated (n=5) and Aldara-treated (n=5) mice. Significantly up- or down-regulated genes are highlighted in bold. **B**, Spontaneous EPSCs on ventral MSNs for control (top) and Aldara-treated mice (bottom) recorded in voltage clamp mode ($V_H$ = -70 mV). Red hatched area in **i** shown in **ii** on an expanded time base. **C & D**, Aldara treatment had no effect on the amplitude or frequency, respectively, of sEPSCs in ventral MSNs: control *vs* Aldara-treated 10.4 ± 0.6 pA *vs* 11.9 ± 0.6 pA; p = 0.1014, Mann-Whitney test (amplitude). Control *vs* Aldara-treated 3.4 ± 0.5 Hz ± 3.5 ± 0.8 Hz; p = 0.6481, Mann-Whitney test (frequency). All data taken from 6 mice per group across 3 different litters, with only 1 neuron taken per brain slice.

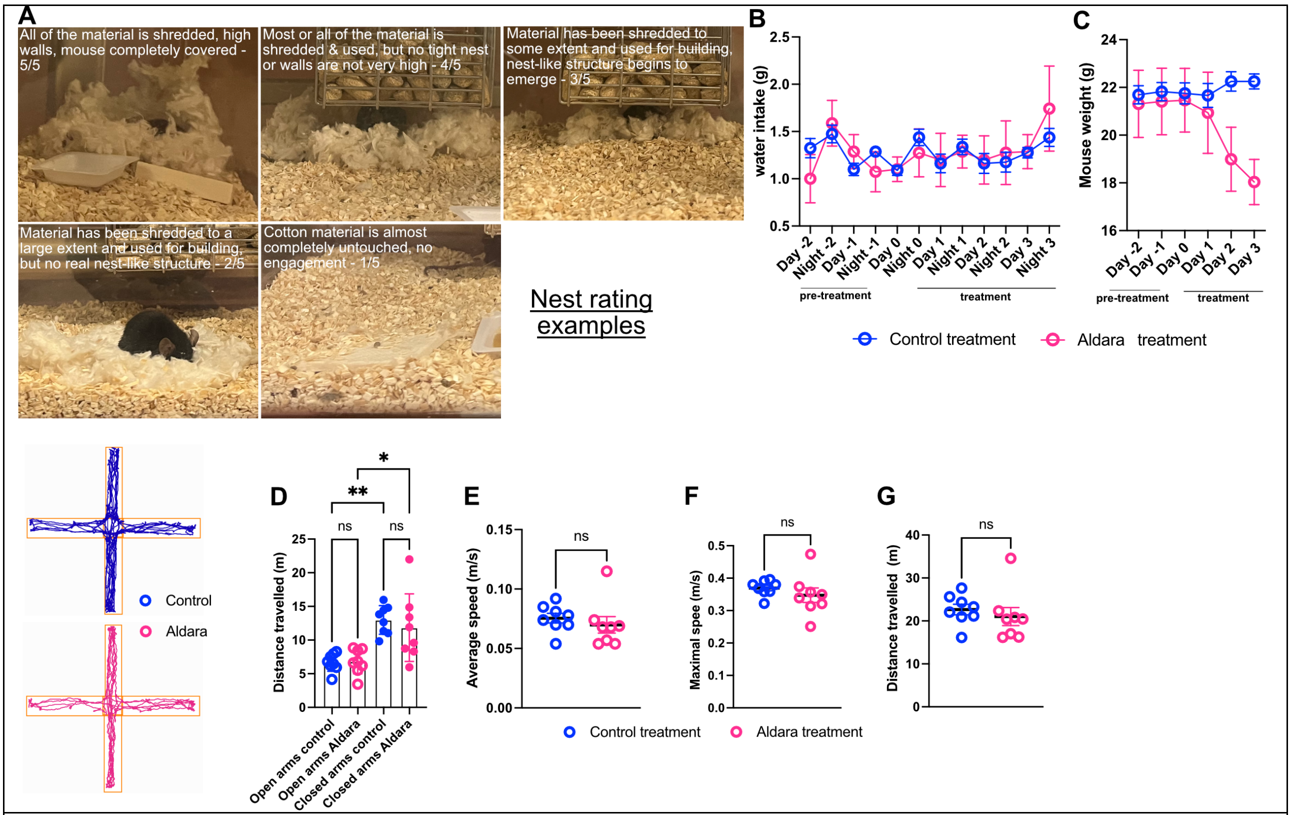


**Supplementary figure 6: Additional behavioural data. A**, example nest ratings from the nest building assay. **B**, while Aldara-treated mice reduced their sucrose consumption relative to control-treated mice, there was no difference in water consumption between the two groups. **C**, Aldara-treated mice displayed significant weight loss over the course of treatment. **D**, there was no difference in anxiety measures (distance travelled in open *vs* closed arms) between Aldara and control mice in the elevated plus maze (EPM) test. **E – G**, Aldara-treated mice displayed no differences in the EPM for average running speed (**E**), maximal running speed (**F**), nor total distance travelled (**G**).

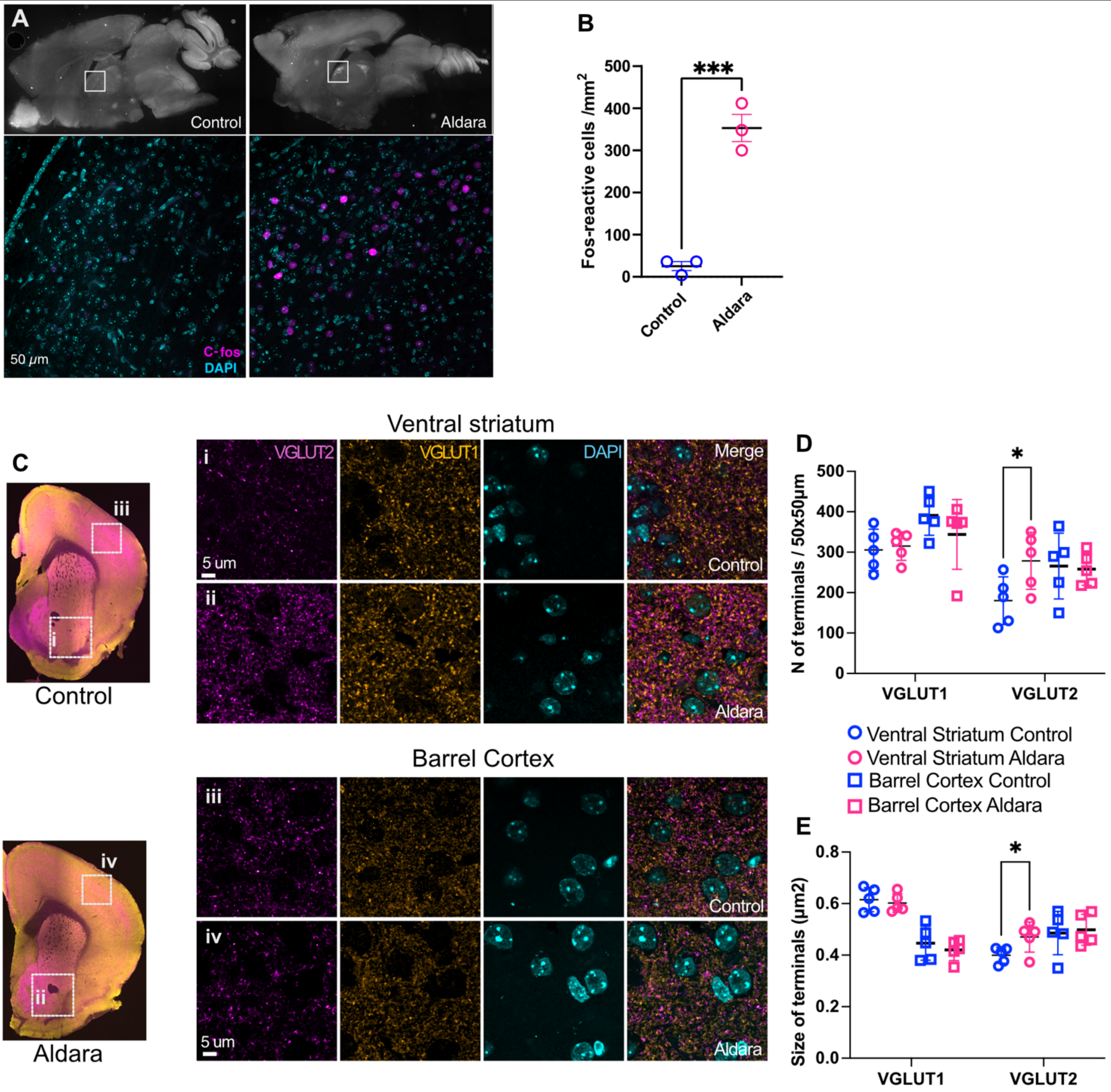


**Supplementary figure 7: Region-specific changes in c-Fos activation and VGlut1 and VGlut2 expression in Aldara-treated mice. A**, Representative images from control and Aldara-treated brain showing fos-positive nuclei (magenta). Scale bar 50 micron. **B**, A significant increase in fos-reactive nuclei in the thalamus 4 hours post treatment, control *vs* Aldara-treated 25.33 ± 18.5 *vs* 353.3 ± 56.2, p = 0.0007, unpaired t-test. **C**, Representative images of VGlut1 and VGlut2 expressing terminals in control (**Ci&Ciii**) and Aldara (**Cii&Civ**) in ventral striatum (**Ci&Cii**) and barrel cortex (**Ciii&Civ**). Scale bar 5 microns. **D**, Number of VGlut2 but not VGlut1 positive terminals in ventral striatum, but not barrel cortex increases in Aldara-treated mice. VGlut1 305.6 ± 51.2 *vs* 315.1 ± 35.2 in ventral striatum; 391.2 ± 39.4 *vs* 343.9 ± 86.2 in barrel cortex; VGlut2 180.0 ± 59.1 *vs* 278.5 ± 70.3 in ventral striatum; p = 0.0251 in *post hoc* Šídák's multiple comparisons test after two-way ANOVA; VGlut2 365.7 ± 81.5 *vs* 258.1 ± 40.0 in barrel cortex. **E**, Size of VGlut2 but not VGlut1 positive terminals in ventral striatum but not barrel cortex increases in Aldara-treated mice. VGlut1 0.615 ±

0.047 *vs* 0.602 ± 0.036 in ventral striatum; 0.446 ± 0.067 *vs* 0.420 ± 0.041 in barrel cortex; VGlut2 0.400 ± 0.031 *vs* 0.470 ± 0.058 in ventral striatum; p = 0.0451 in *post hoc* Šídák's multiple comparisons test after two-way ANOVA; VGlut2 0.486 ± 0.084 *vs* 0.498 ± 0.060 in barrel cortex.

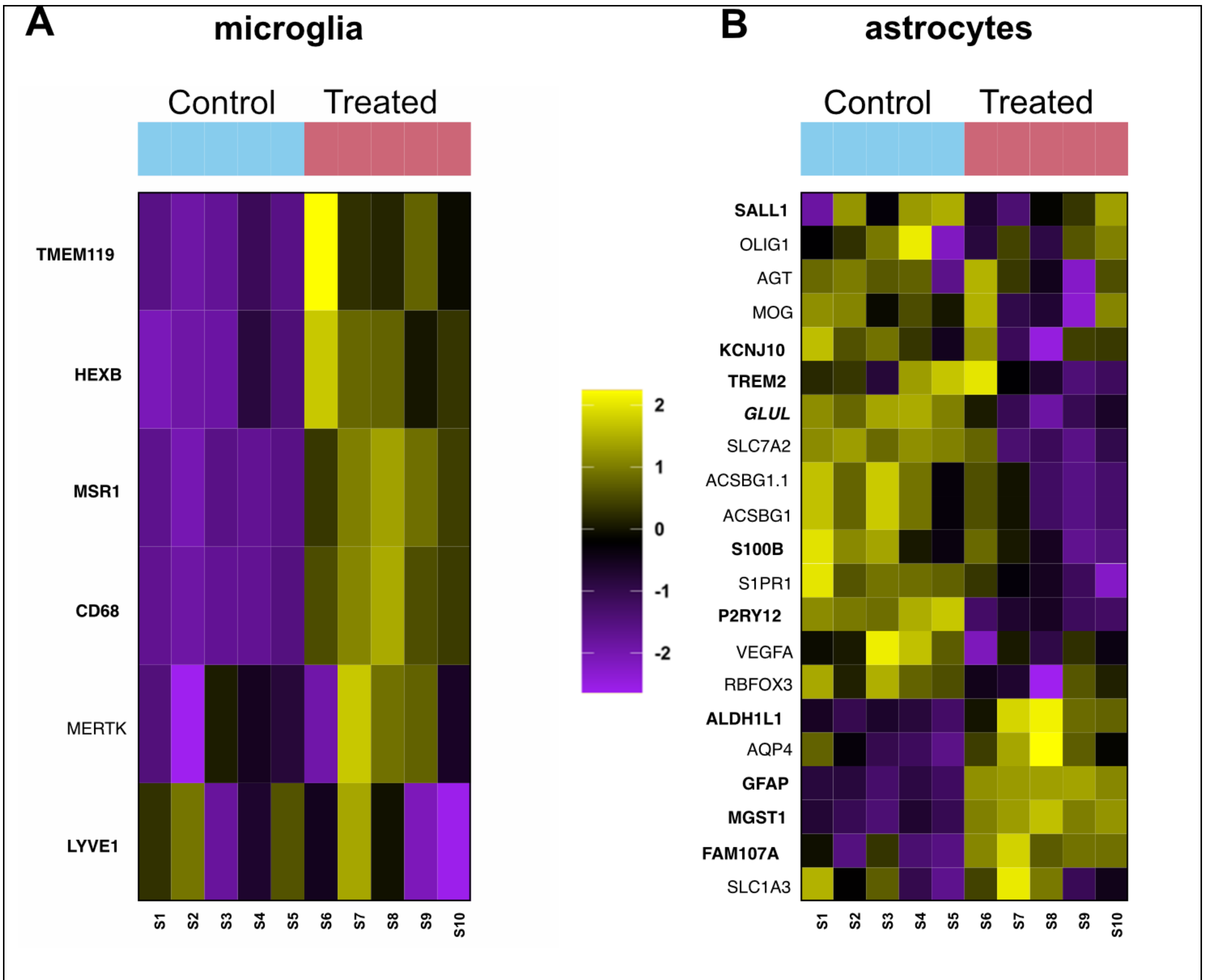


**Supplementary figure 8: Differential expression analysis for microglia and astrocyte-related genes.** Heatmaps comparing expression levels for selected genes between control- (left) and Aldara-treated mice for genes associated with **A**, microglial or **B**, astrocytic activation. Genes highlighted in bold were displayed significant differences in expression levels between the 2 groups. Each block represents one mouse (n=5 per group).

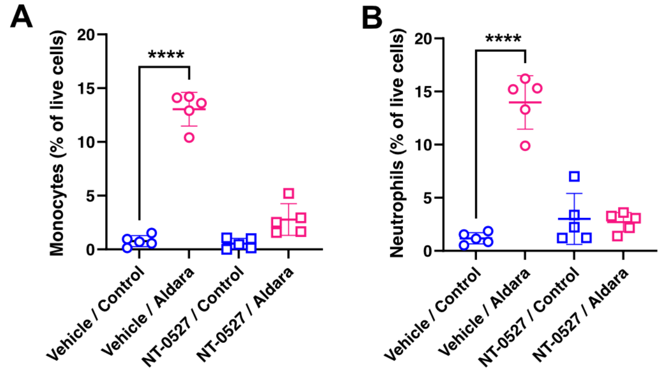


**Supplementary figure 9: Treatment with the NLRP3 inhibitor, NT-0527, blocks Aldara-mediated increase in peripheral monocytes and neutrophils. A**, Flow cytometry counts for total number of monocytes as a percentage of live cells. **B**, flow cytometry counts for total number of neutrophils as a percentage of live cells. ****, p< 0.0001, *post hoc* Sídák's multiple comparisons test after ANOVA.

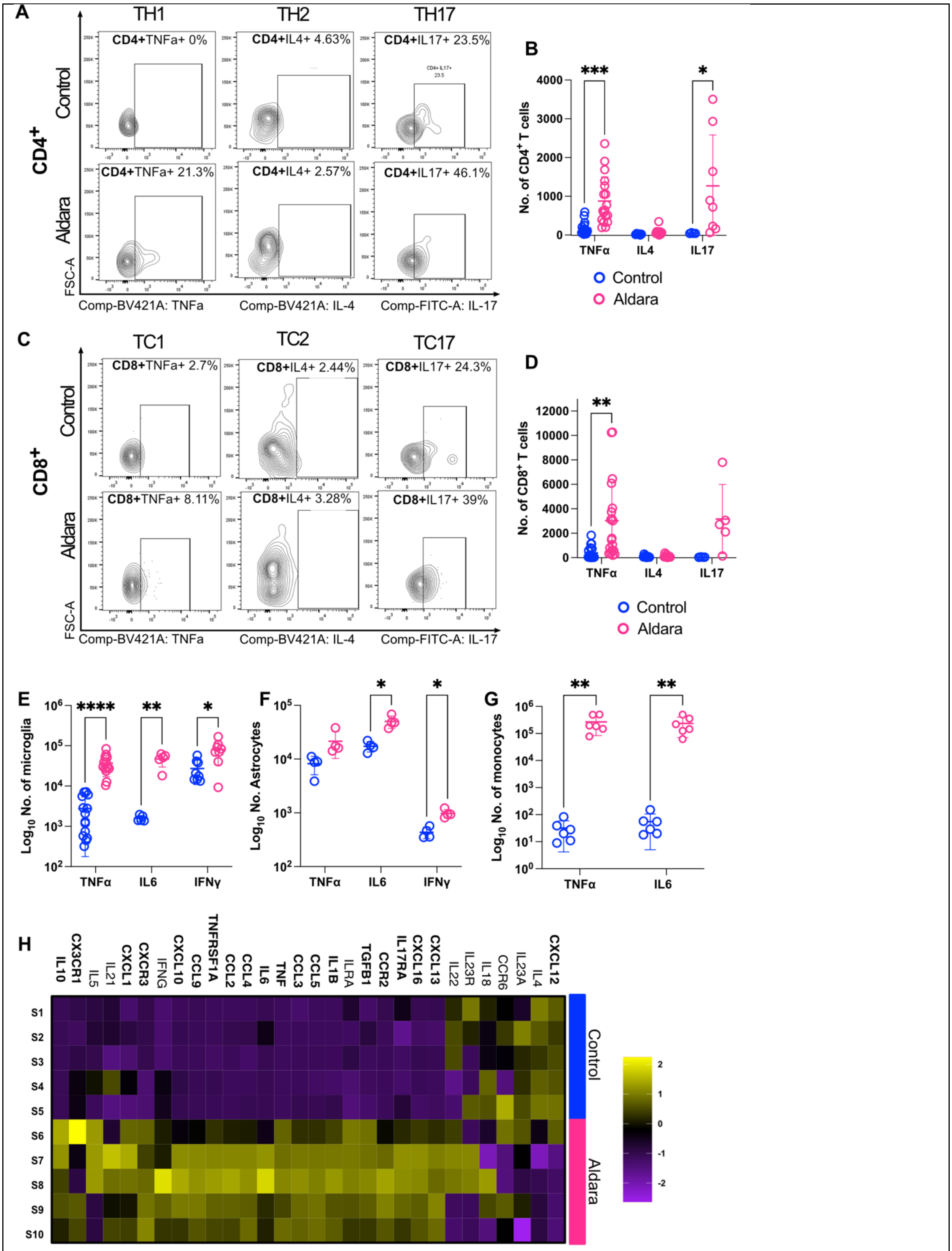


**Supplementary figure 10: Aldara induces a neuroinflammatory response via multiple T cell subtypes.** In all panels control mice are shown in blue and Aldara-treated mice in pink. **A&C**, Contour maps showing the numbers of CD4$^+$ (upper two rows) and CD8$^+$ (lower two rows) T cells in brain homogenates from control- and Aldara-treated mice. **B**, Expression of

Th1 type cytokine TNFα and IL17 increases in Aldara-treated brains **but no** change in IL4, a Th2-type cytokine, was observed in $CD4^+$ T cells as compared to control; TNFα n= 174 ± 174 *vs* 876 ± 625, p = 0.0006 , IL-4 n= 20 ± 13 *vs* 88 ± 102 , IL-17 n= 47 ± 11 *vs* 1268 ± 1314.2, as found with *post hoc* Šídák's multiple comparisons test after mixed-effects analysis. **D**, Aldara treatment resulted in an increase in TNFα expression in $CD8^+$ T cells as compared to control. TNFα n= 389 ± 530 *vs* 3022 ± 3183, p = 0.0063, IL-4 n= 99 ± 85 *vs* 130 ± 114, IL17 n= 39 ± 14 *vs* 3127 ± 2827, as found with *post hoc* Šídák's multiple comparisons test after mixed-effects analysis. **E**, Microglia displayed a significant increase in all three cytokines, as compared to control, TNFα n= 2764 ± 2587 *vs* 37010 ± 19800, p = < 0.0001 (TNFα), IL6 n= 1590 ± 265 *vs* 46799 ± 17232, p = 0.0126 (IL6), IFN-γ n= 27259 ± 16448 *vs* 80073 ± 46930, p = 0.0456 as found with *post hoc* Šídák's multiple comparisons test after mixed-effects analysis. **F**, Astrocytes displayed no change in TNFα but a significant increase in IL6 and IFN-γ, TNFα n= 8176 ± 3071 *vs* 21457 ± 11197, IL6 n= 17361 ± 3795 *vs* 50294 ± 12936, p = 0.0334, IFN-γ n= 435 ± 105 *vs* 976 ± 176, p = 0.0102 as found with *post hoc* Šídák's multiple comparisons test after mixed-effects analysis. **G**, Infiltrating peripheral myeloid cells showed a significant increase in TNFα and IL6 expression as compared to control, TNFα n= 32 ± 28 *vs* 267187 ± 183460, p = 0.0025, IL6 n= 55 ± 50 *vs* 235076 ± 164587, p = 0.0071 as found with *post hoc* Šídák's multiple comparisons test after mixed-effects analysis. Data plotted as mean ± SD. **H**, Searchlight heatmap showing relative whole brain expression levels for a range of cytokines from control- and Aldara-treated mice. Each square on the heatmap represents one mouse (S1 to S10).

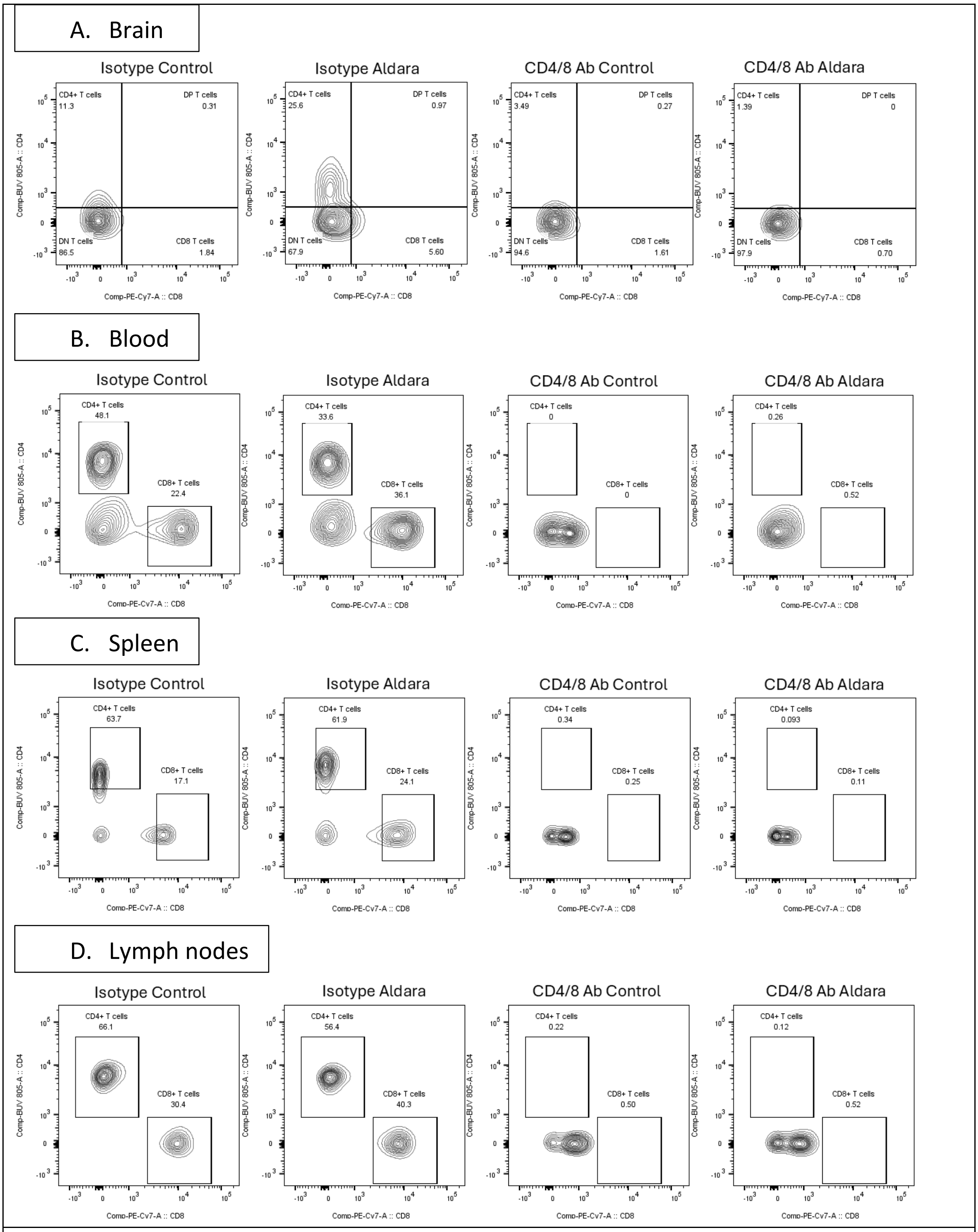


**Supplementary figure 11: Whole body CD4 and CD8 depletion using antibodies.** CD4 and CD8 contour flow plots of single cell homogenates from Brain (A), Blood (B), Spleen (C) and Lymph nodes (D) from mice injected with 400µg of rat IgG isotype (clone LTF-2), 200µg CD4 (clone GK1.5) and 200µg CD8 (clone 2.43) antibodies, intra peritoneum (*ip*), three days before starting the control and Aldara cream treatment and again couple of hours before the 2nd day of cream treatment. Tissue collection was done after final (3rd day) of cream application for flow cytometry.

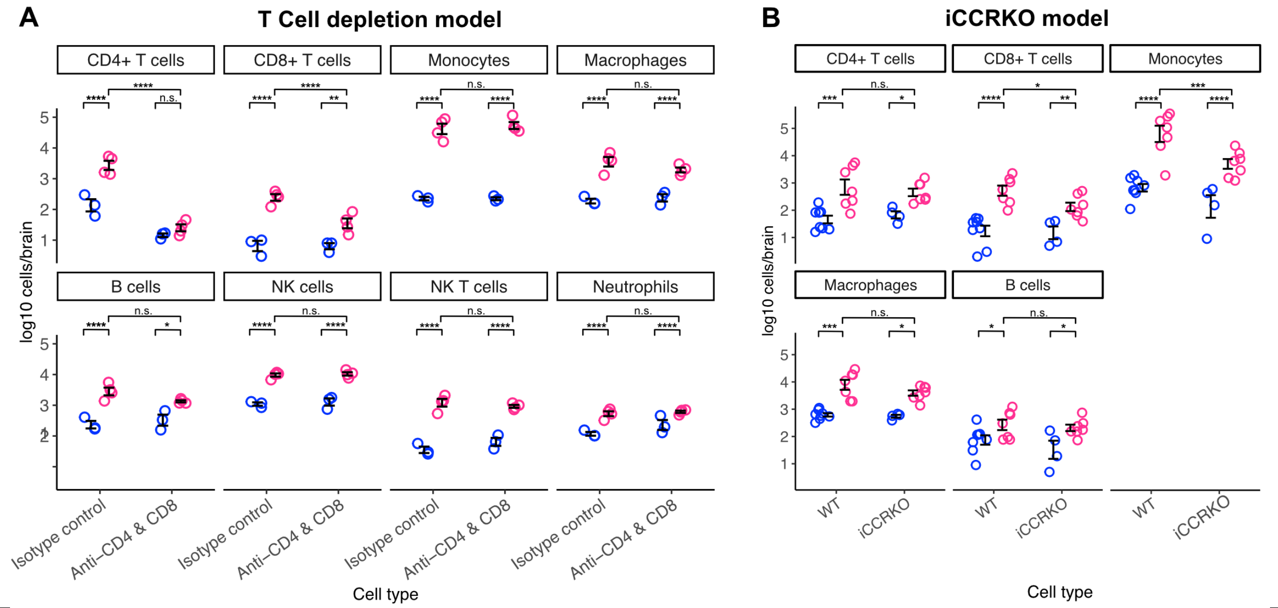


**Supplementary figure 12: Control data for T cell depletion and iCCR$^{-/-}$ experiments.** Flow cytometry cell counts. **A,** in the T cell depletion model, we saw a complete inhibition of CD4+ T cell ingress into the brain after Aldara treatment, and a substantial attenuation of the CD8+ T cell response. Importantly, we saw no effect of T cell depletion on the ingress of other immune cells, notably monocytes, into the brain after Aldara treatment. Overall, we found a significant effect of Aldara treatment ($p<1x10^{-9}$) and interaction between Aldara and depletion status (p=0.0000374) using a linear mixed effects model of $log_{10}$-transformed data, modelled in R. **B,** similarly, in the iCCR$^{-/-}$ mouse, we saw a significant attenuation of infiltrating monocytes into the brain of iCCR$^{-/-}$ compared with wildtypes after Aldara treatment. We saw no effect of chemokine receptor deletion on the ingress of other immune cells in response to Aldara treatment. Overall, we found a significant effect of Aldara treatment (p=0.0000231) and no overall interaction between Aldara treatment and genotype (p=0.3671), but a significant interaction between genotype and monocyte ingress (p=0.00223), using a linear mixed effects model of $log_{10}$-transformed data, modelled in R. *, p<0.05; **, p<0.01; ***, p<0.001; ****, p<0.0001. All post-hoc tests were correct for multiple comparisons across the entire flow experiment using the FDR method. Linear mixed effects models treated individual animals as a random effect, which had a significant effect in both experiments.

### Supplementary Tables

#### Supplementary Table 1: Clinical features of PsD cohort.

| **Clinical features** (n=46) | |
|---|---|
| Age (mean ± SD) | 49 ± 11.2 |
| Disease duration (years, mean ± SD) | 6.4 ± 5.8 |
| Sex (male/female) | 22/24 |
| BMI (mean ± SD) | 29 ± 4.5 |
| Participant gVAS (mean ± SD) | 59 ± 22 |
| DAPSA (mean ± SD) | 40.8 ± 19 |
| Fatigue NRS (mean ± SD) | 6.5 ± 2.6 |
| PROMIS Fatigue (mean ± SD) | 51.3 ± 16.6 |
| PROMIS Fatigue motivational impact (mean ± SD) | 13.4 ± 4.2 |
| PROMIS Fatigue motivation M3* | 3.3 ± 1 |
| PROMIS Fatigue motivation M4† | 3.7 ± 1 |
| PROMIS Depression (mean ± SD) | 19.1 ± 9.8 |
| PROMIS Depression severity (%) | |
| • None-slight | 41.3% |
| • Mild | 30.4% |
| • Moderate | 15.2% |
| • Severe | 13% |

BMI – Body Mass Index; DAPSA – Disease Activity in PSoriatic Arthritis; gVAS – Global Disease Activity; NRS – numeric rating scale; PROMIS – Patient-Reported Outcomes Measurement Information System.

* M3: How often were you less effective at home due to fatigue?
† M4: How often did you have to push yourself to get things done because of your fatigue?
