## Supplementary methods for "Psoriasis-related neuroinflammation disrupts thalamostriatal signalling driving anhedonia in both humans and mice"

1. Master Gating strategy

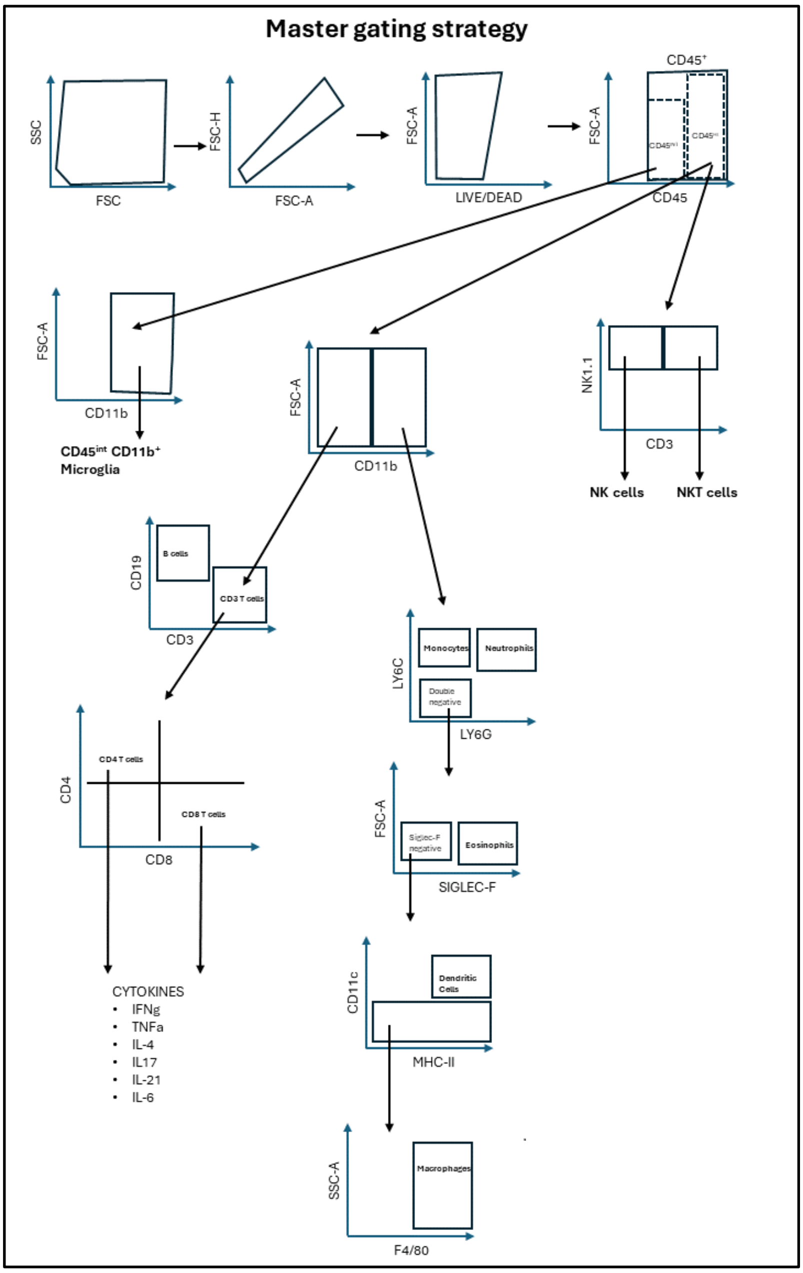

1. Gating strategy for basic 4 population

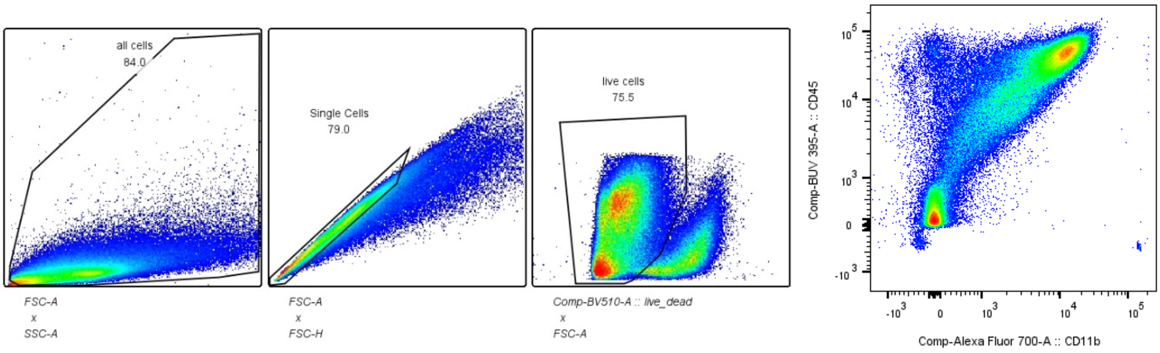

2. Immune cell (CD45+) gating strategy with FMO

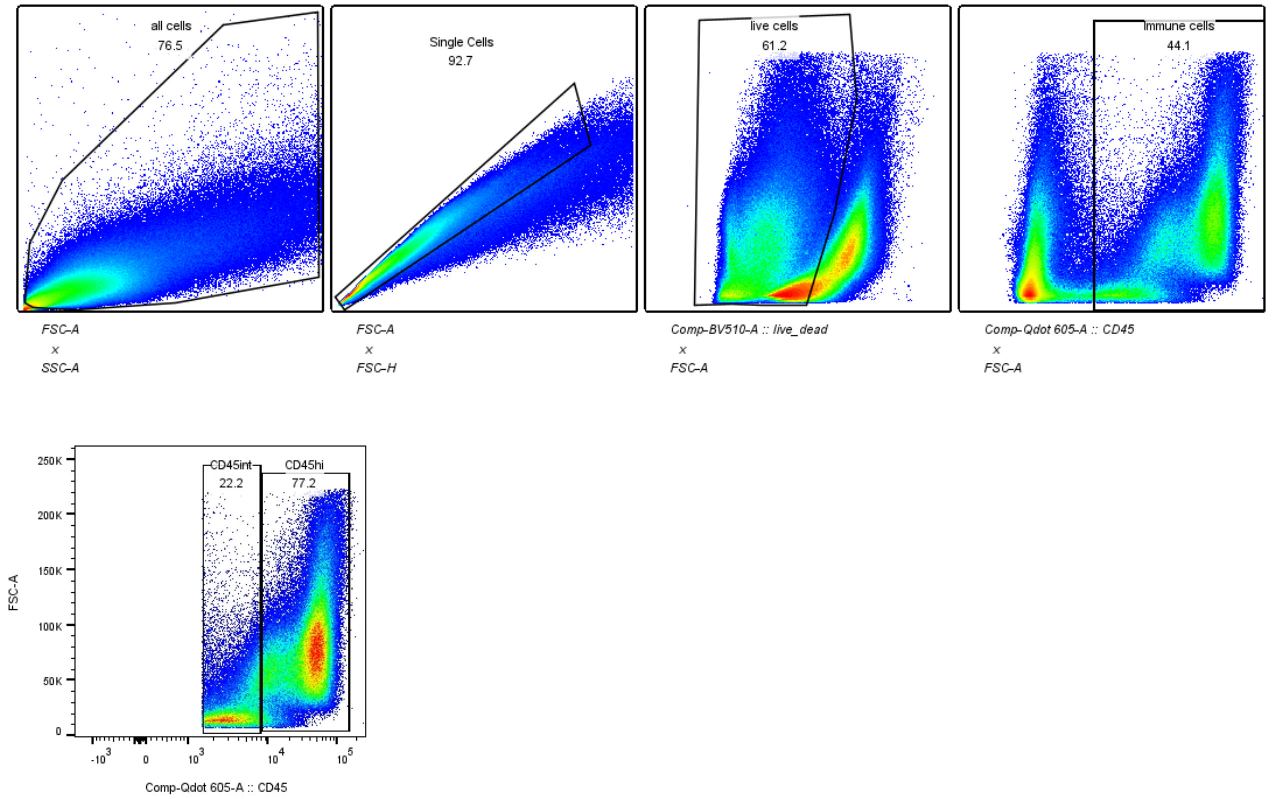

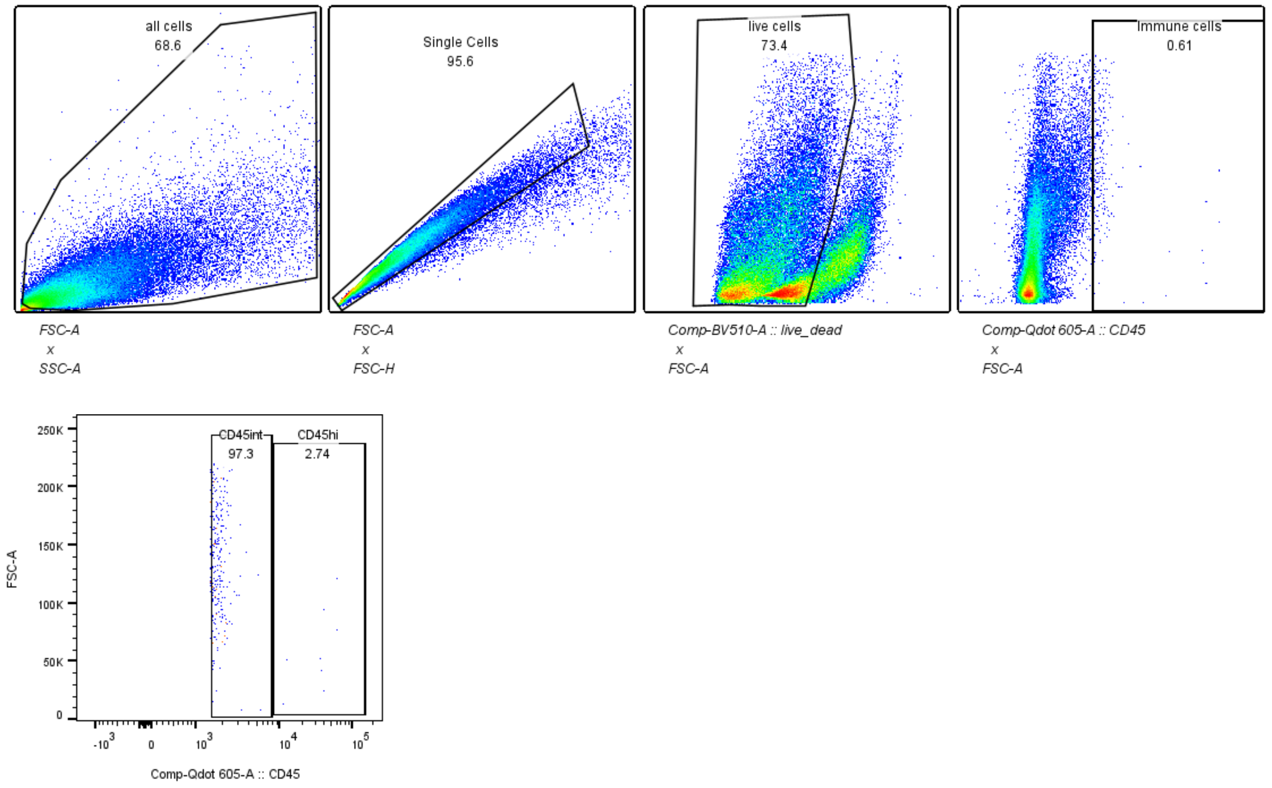

### 3. Monocyte gating with FMO

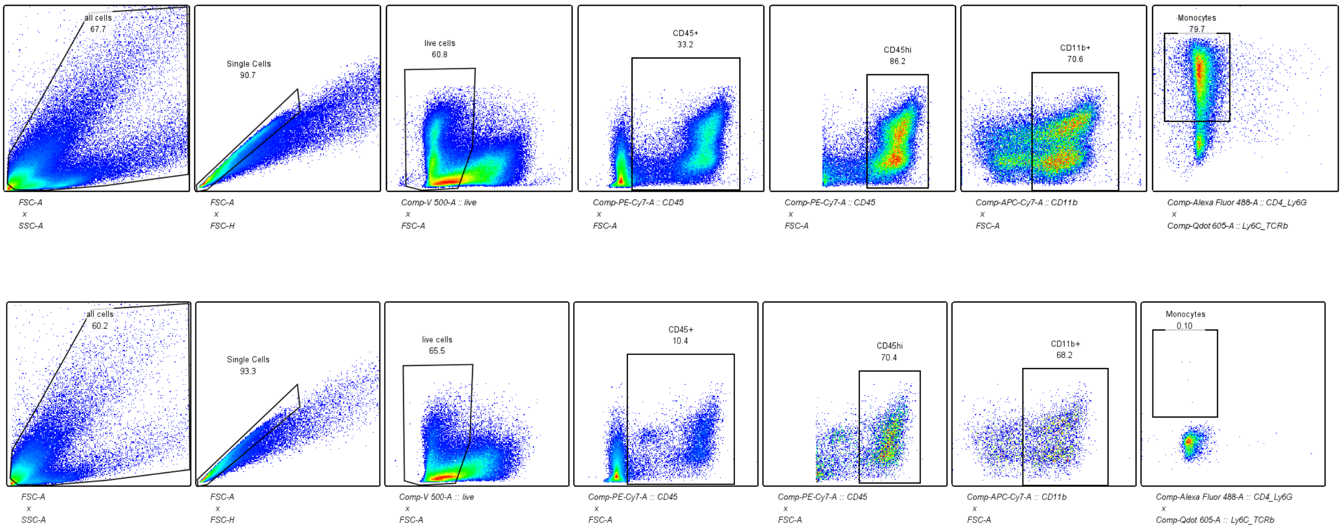

### 4. B cell gating strategy and FMO

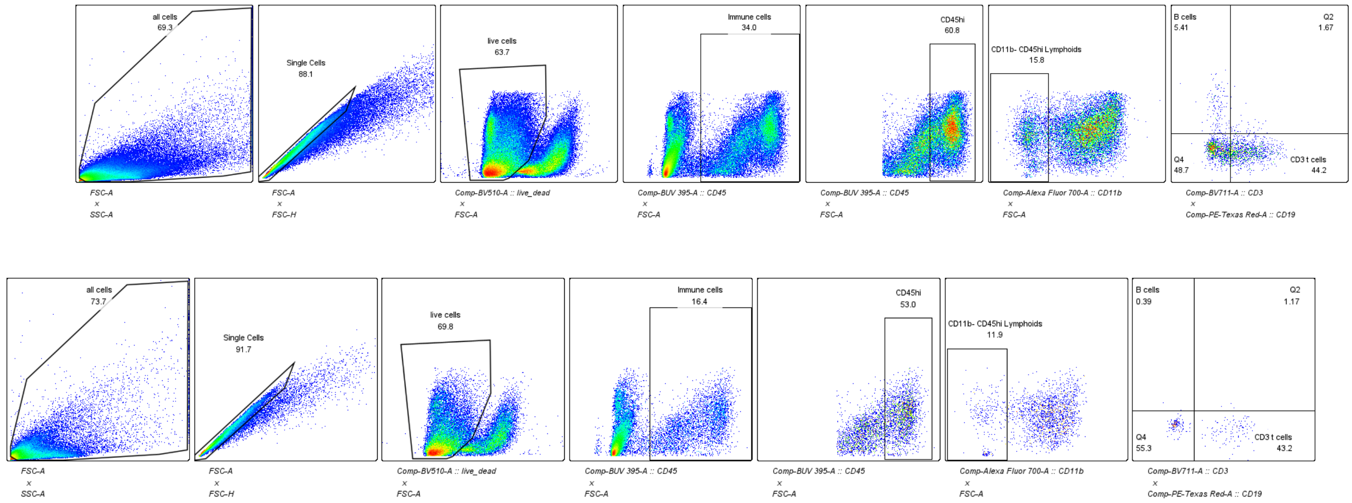

5. NK cell gating strategy and FMO

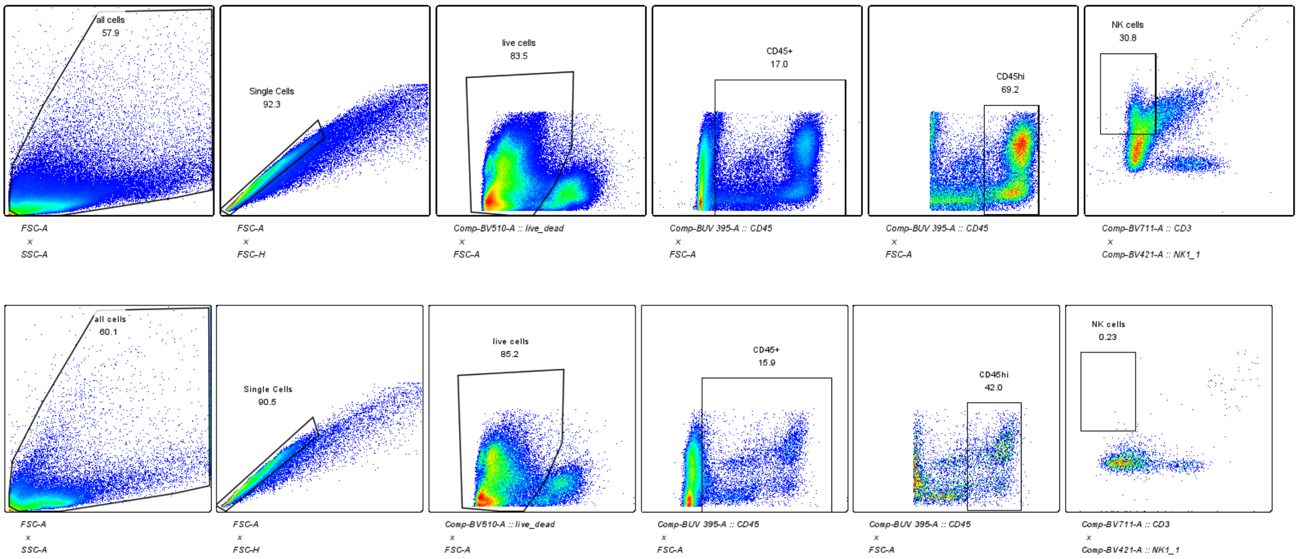

6. NKT cells gating strategy and FMO

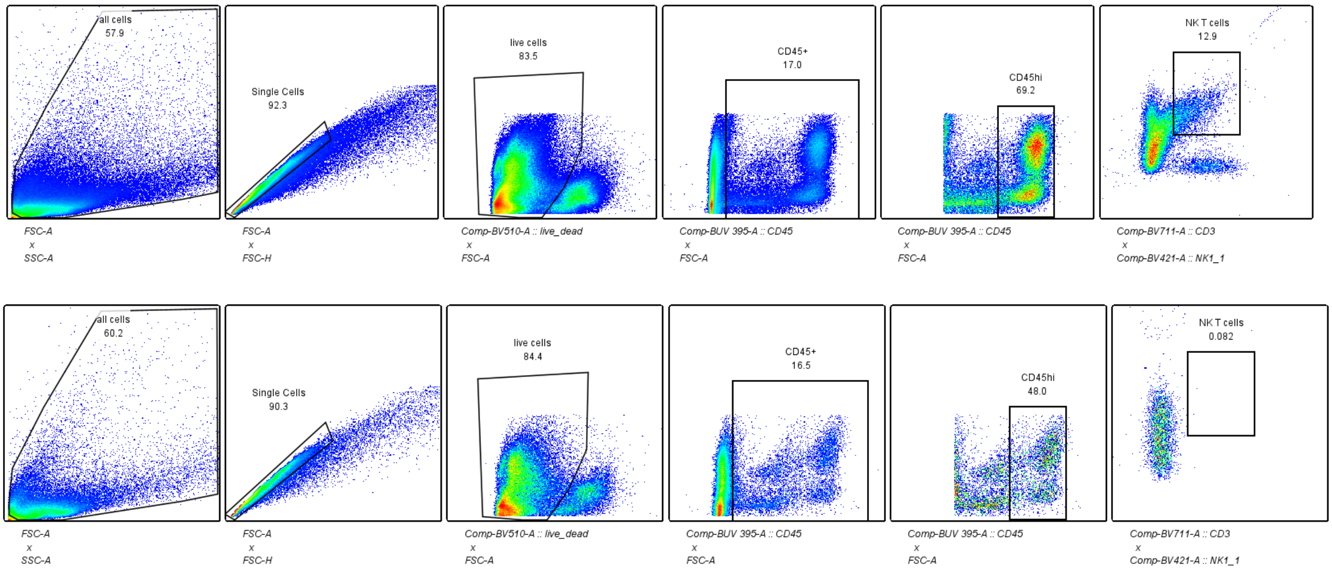

7. CD3 T cell gating and FMO

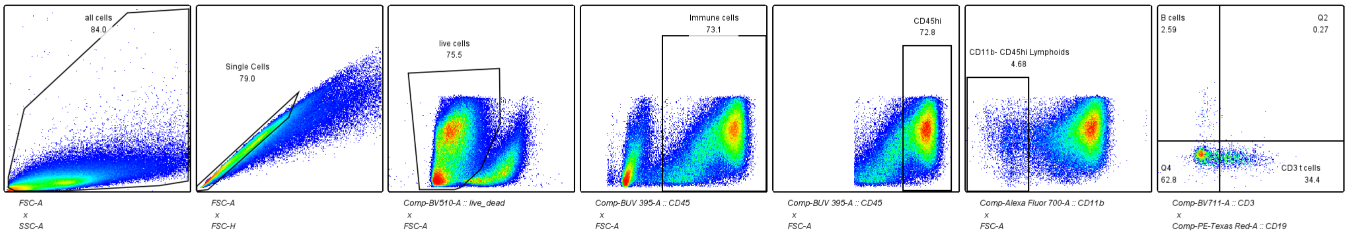

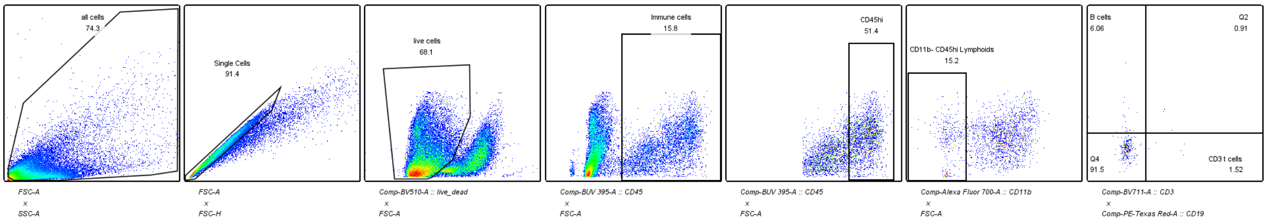

### 8. CD4 gating and FMO

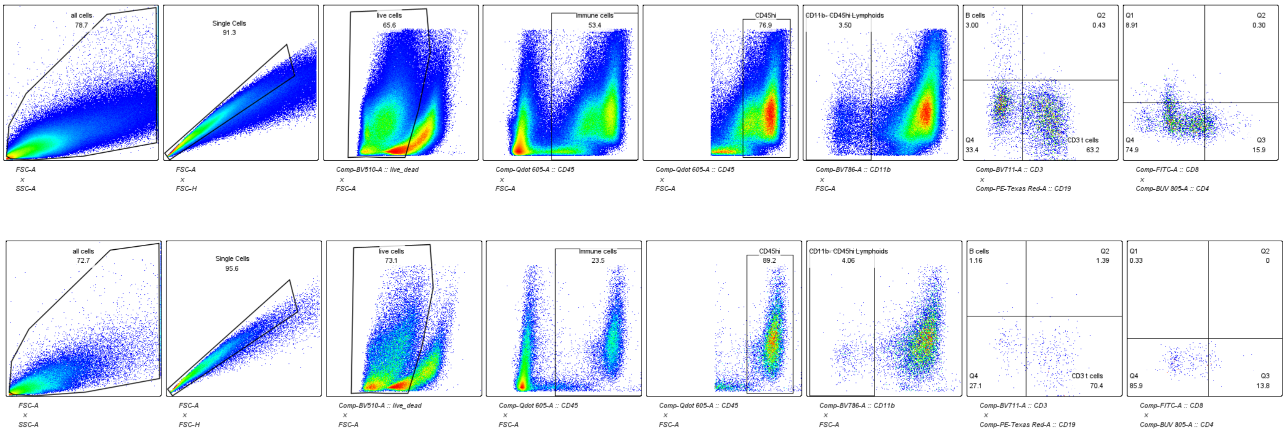

### 9. CD8 gating and FMO

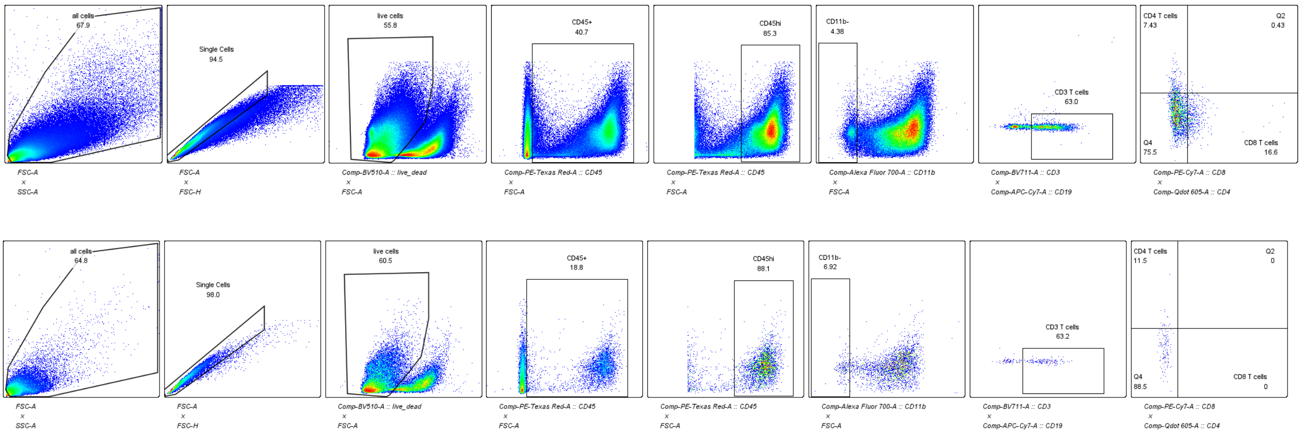

### 10. Microglia gating and FMO

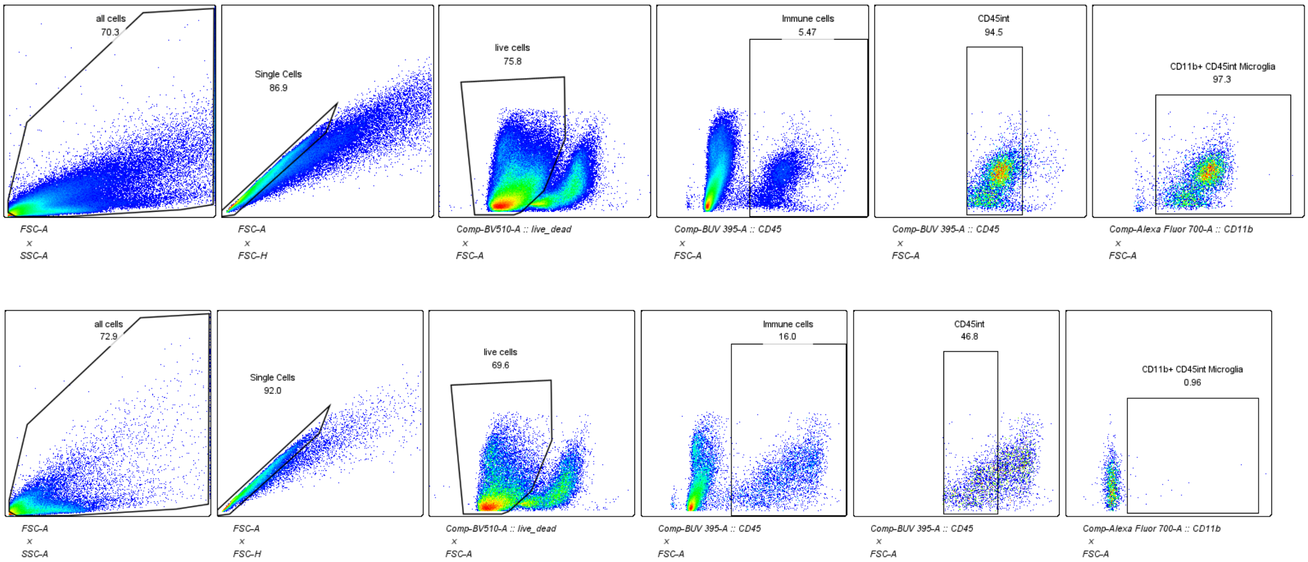

### 11. Astrocytes and their FMOs

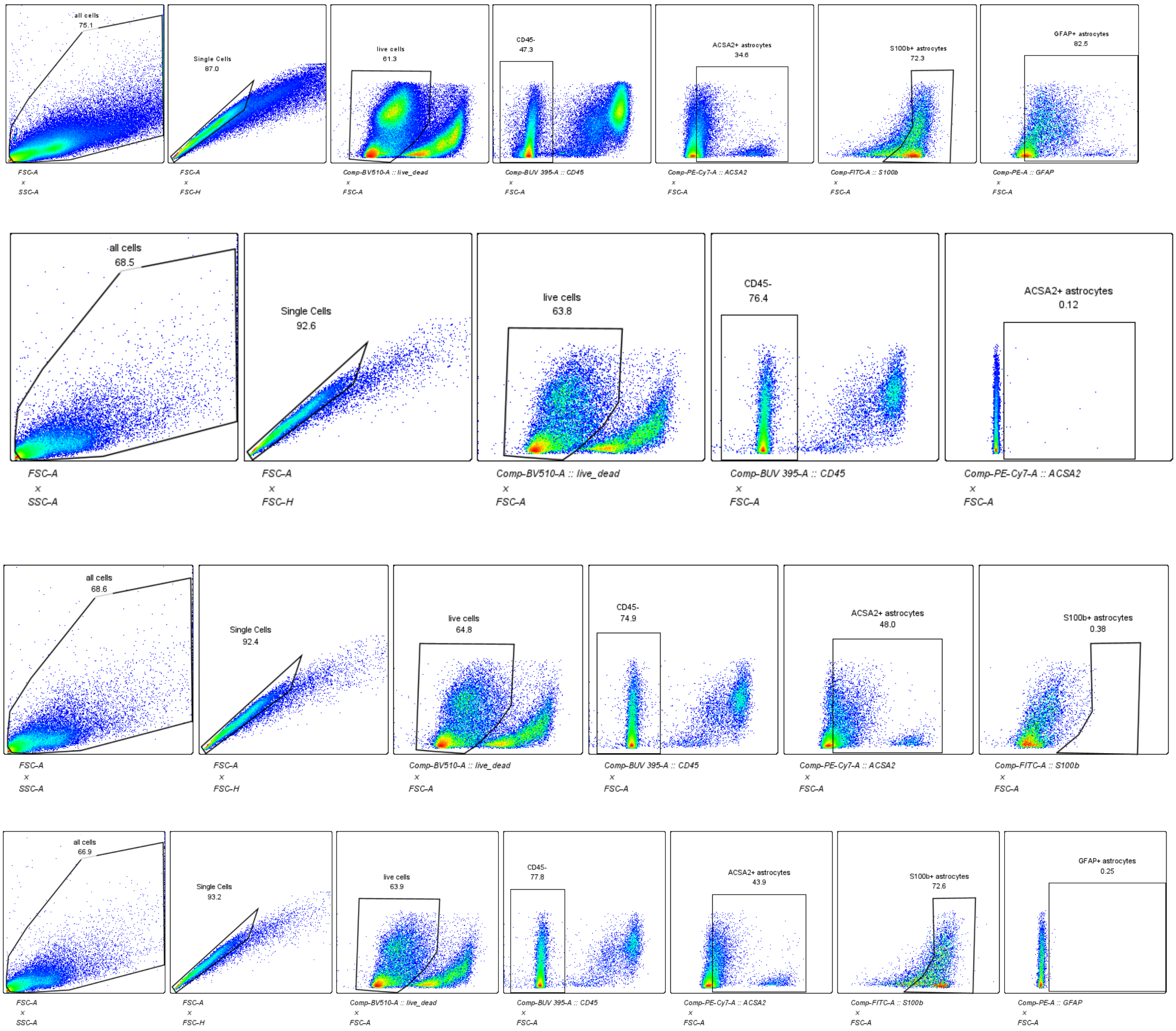

CYTOKINES

1. Microglia TNFa+gating and FMO

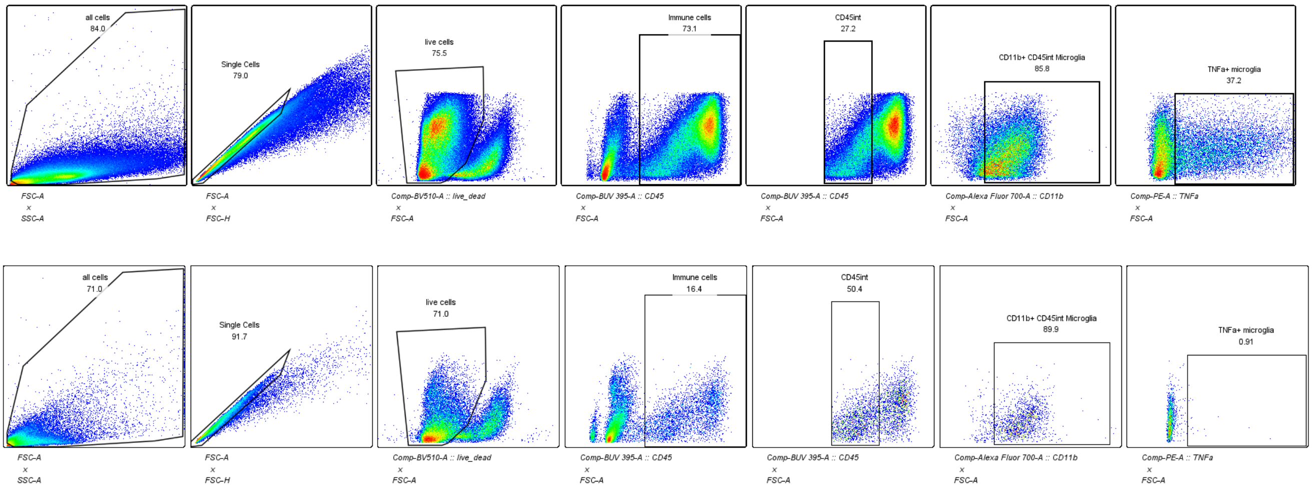

2. CD4+ TNFa+ gating FMO

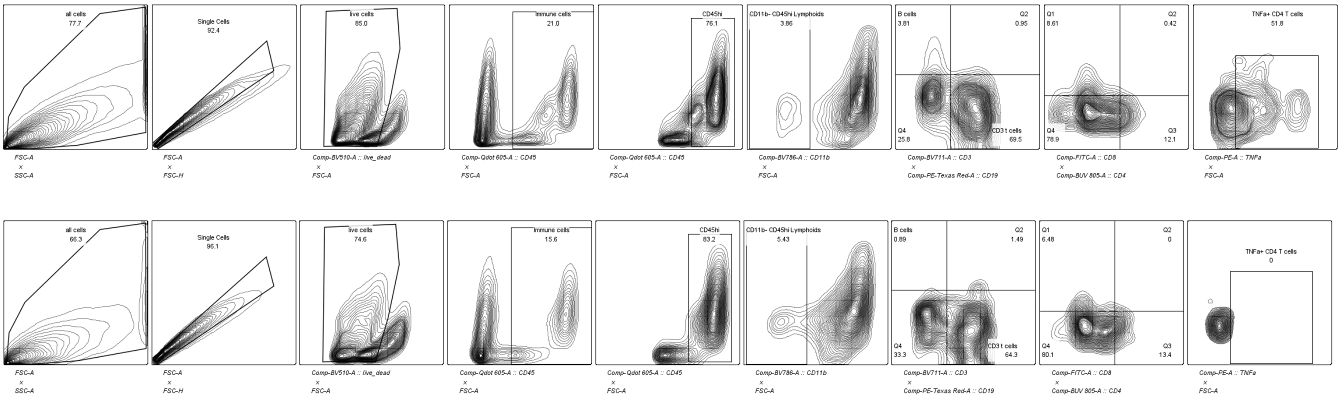

3. CD4+ IL4+ gating FMO

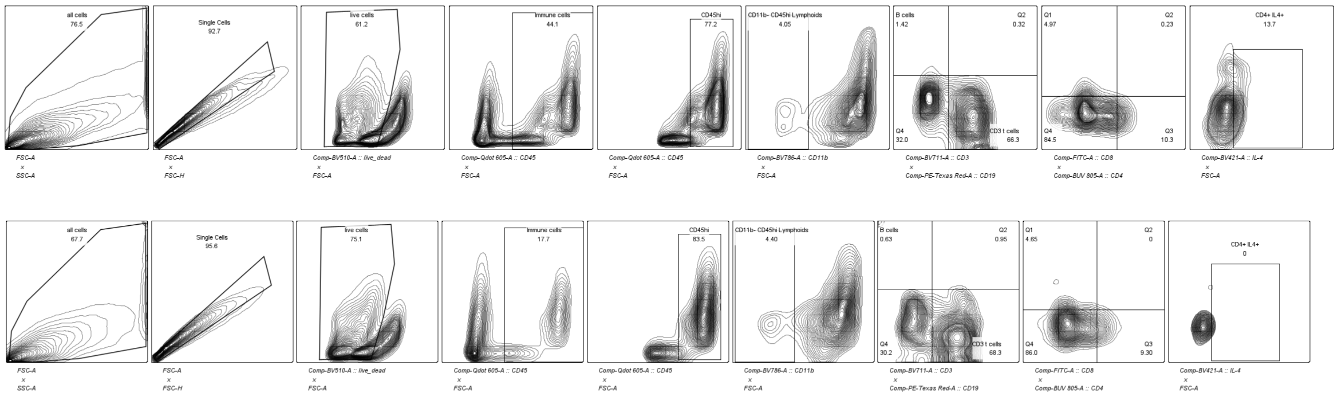

4. CD8+ IL17+ gating and FMO

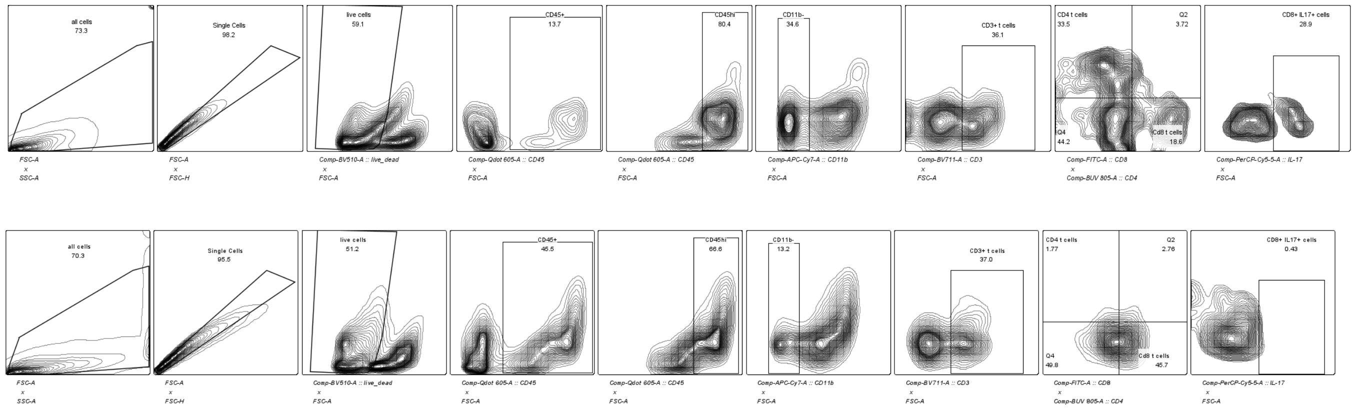

5. CD4+ IL17+ gating and FMO

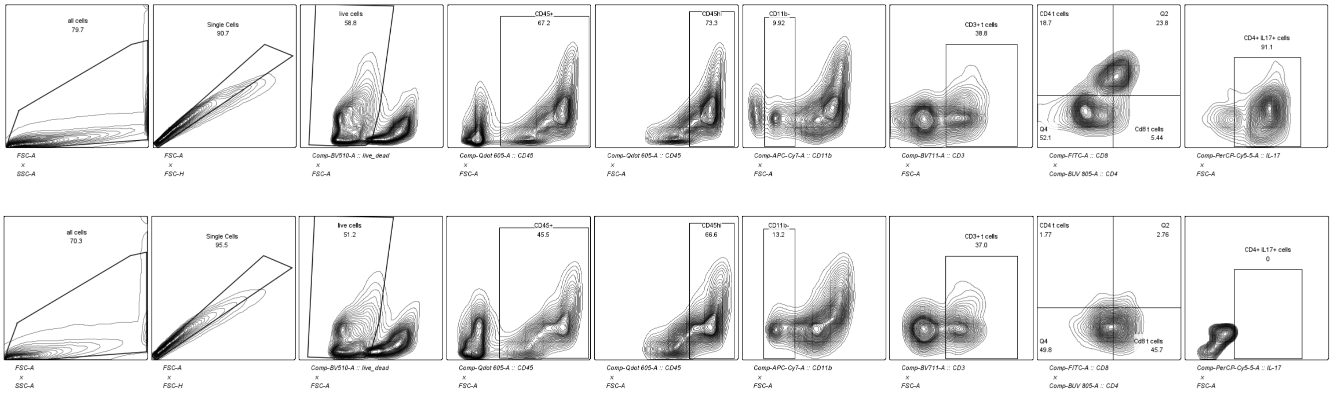

6. CD8+ IL4+ gating FMO

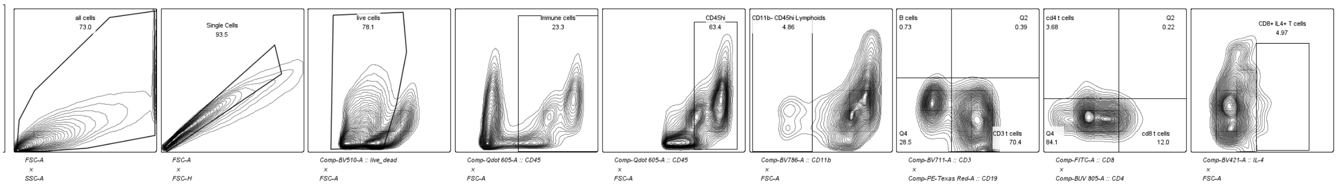

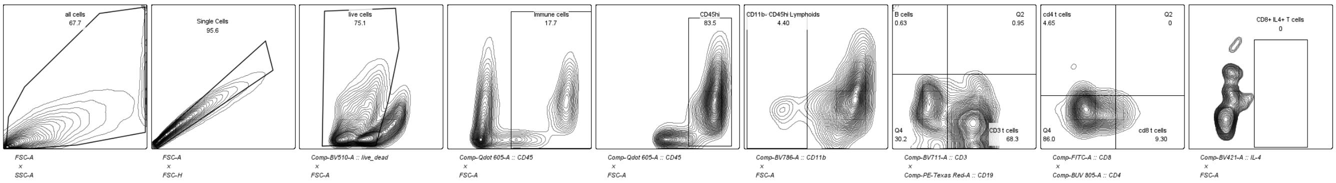

7. CD8+ TNFa+ gating FMO

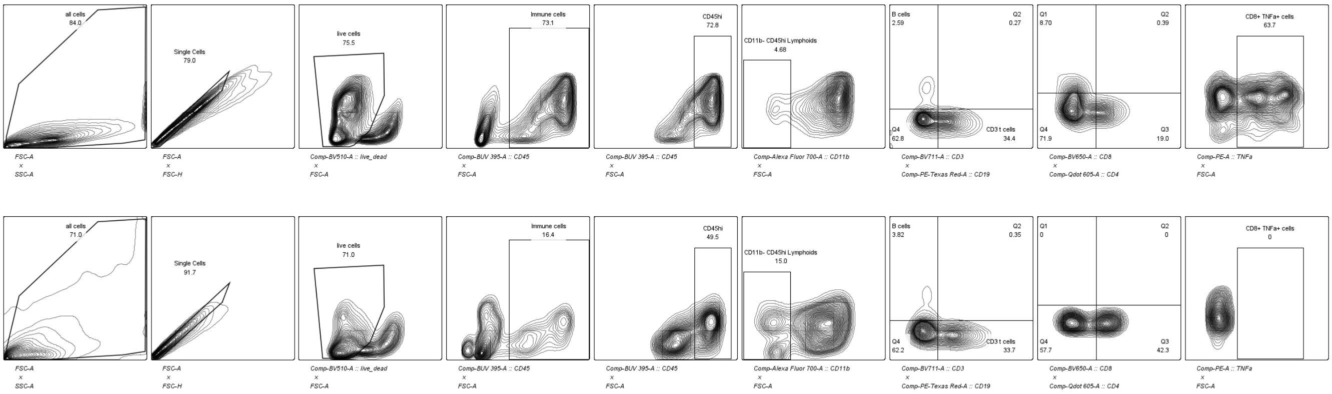

### Supplementary table 2: antibody list for flow cytometry experiments

| No | Marker | Fluorophore | Company | Catalogue No | Clone | RRID | Dilutions |
|---|---|---|---|---|---|---|---|
| 1 | CD45 | PE Texas Red | Biolegend | 103146 | 30-F11 | AB_2564002 | 1:500 |
| | | PCP Cy5.5 | Biolegend | 103132 | 30-F12 | AB_893340 | 1:500 |
| | | BUV395 | BD Biosciences | 564279 | 30-F11 | AB_2651134 | 1:500 |
| | | Bv605 | Biolegend | 103139 | 30-F11 | AB_2562341 | 1:500 |
| 2 | CD11b | AF700 | Biolegend | 101222 | M1/70 | AB_493705 | 1:500 |
| | | APCCy7 | Biolegend | 101226 | M1/70 | AB_830642 | 1:500 |
| 3 | CD3 | Bv711 | Biolegend | 100241 | 17A2 | AB_2563945 | 1:500 |
| | | FITC | Biolegend | 100203 | 17A2 | AB_312660 | 1:500 |
| 4 | CD4 | Bv605 | Biolegend | 100451 | GK1.5 | AB_2564591 | 1:500 |
| | | AF488 | Biolegend | 100406 | GK1.5 | AB_312691 | 1:500 |
| 5 | CD8 | PECy7 | Biolegend | 100722 | 53-6.7 | AB_312761 | 1:500 |
| | | FITC | Biolegend | 100723 | 53-6.7 | AB_389304 | 1:500 |
| 6 | NK1.1 | PCP Cy5.5 | Biolegend | 108916 | DX5 | AB_2129358 | 1:500 |
| 7 | CD19 | PE Texas Red | Biolegend | 115554 | 6D5 | AB_2564001 | 1:500 |
| 8 | Ly6C | APC | Biolegend | 128016 | HK1.4 | AB_1732076 | 1:500 |
| | | AF700 | Biolegend | 128024 | HK1.4 | AB_10643270 | 1:500 |
| 9 | Ly6G | Bv650 | Biolegend | 127641 | 1A8 | AB_2565881 | 1:500 |
| | | AF488 | Biolegend | 127606 | 1A8 | AB_1236494 | 1:500 |
| | | BUV395 | BD Biosciences | 563978 | 1A8 | AB_2716852 | 1:500 |
| 11 | SiglecF | PE | BD Biosciences | 552126 | E50-2440 | AB_394341 | 1:500 |
| 12 | CD11c | APCCy7 | Biolegend | 117323 | N418 | AB_830646 | 1:500 |
| | | AF700 | Biolegend | 117320 | N418 | AB_528736 | 1:500 |
| 13 | MHCII | Bv786 | Biolegend | 107645 | M5/114.15.2 | AB_2565977 | 1:500 |
| | | FITC | Biolegend | 107616 | M5/114.15.2 | AB_493523 | 1:500 |
| | | Bv605 | Biolegend | 107639 | M5/114.15.2 | AB_2565894 | 1:500 |
| 14 | F4/80 | BUV805 | BD Biosciences | 749282 | T45-2342 | AB_2873657 | 1:500 |
| 17 | MSR1 | BV711 | BD Biosciences | 268318 | Clone 268318 | AB_2872549 | 1:200 |
| 18 | MERTK | PE Texas Red | biolegend | 151524 | 2B10C42 | AB_2876509 | 1:500 |
| 19 | GFAP | FITC | biolegend | 644704 | 2E1.E9 | AB_2566109 | 1:100 |
| 20 | O4 | APC | Miltenyi Biotec | 130-119-155 | O4 | AB_2751912 | 1:100 |
| 21 | ASCA2 | PE CY7 | Miltenyi Biotec | 130-116-246 | REA969 | AB_2727424 | 1:100 |
| | S100beta | FITC | Biotechne | NBP2-59619F | | | 1:100 |
| | GFAP | PE | Miltenyi Biotec | 130-118-351 | REA335 | AB_2651834 | 1:100 |
| 22 | CD24 | Bv605 | biolegend | 101827 | M1/69 | AB_2563464 | 1:500 |
| 23 | CD29 | AF700 | biolegend | 102218 | HMb1-1 | AB_493711 | 1:500 |
| 25 | IL17 | AF488 | biolegend | 506910 | TC11-18H10.1 | AB_536012 | 1:100 |
| 26 | IFNg | APC | biolegend | 505810 | XMG1.2 | AB_315404 | 1:100 |
| | | Bv421 | biolegend | 505830 | XMG1.2 | AB_2563105 | 1:100 |
| 27 | IL4 | Bv421 | biolegend | 504119 | 11B11 | AB_10896945 | 1:100 |
| 28 | TNFa | PE | BD Biosciences | 554419 | MP6-XT22 (RUO) | AB_395380 | 1:100 |
| | | Bv421 | biolegend | 506327 | MP6-XT22 | AB_10900823 | 1:100 |
| 29 | IL6 | APC | biolegend | 504508 | MP5-20F3 | AB_10694868 | 1:100 |

| | | | | | | | |
|---|---|---|---|---|---|---|---|
| 30 | CD4 | | BioXcell (invivomab) | BE0003-1 | GK1.5 | AB_1107636 | |
| 31 | Isotype | IgG2b | BioXcell (invivomab) | BE0090 | | AB_1107780 | |
| 32 | CD8 | | BioXcell (invivomab) | BE0004-1 | 53-6.7 | AB_1107671 | |
| 33 | Isotype | IgG2a | BioXcell (invivomab) | BP0089 | | AB_1107769 | |
| 35 | Foxp3 | AF700 | biolegend | 126421 | MF-14 | AB_2750492 | 1:100 |
| 32 | CD103 | APC | biolegend | 110905 | W19396D | AB_2927988 | 1:500 |
| 34 | CTLA4 | PE-Cy7 | biolegend | 106313 | UC10-4B9 | AB_2564237 | 1:500 |
| 33 | CD69 | Bv650 | biolegend | 104541 | H1.2F3 | AB_2616934 | 1:500 |

### Supplementary table 3: antibody list for immunoblotting experiment

| S. No. | Marker | Fluorophore | Company | Catalogue No | Clone | RRID | Dilutions |
|---|---|---|---|---|---|---|---|
| 1 | NLRP3 | | Cell Signalling | 15101 | D4D8T | | 1:1000 |
| 2 | IL1b | | Cell Signalling | 12242S | 3A6 | | 1:1000 |
| 3 | Goat antimouse IgG | HRP | ThermoScientific | 31430 | | AB_228307 | 1:10000 |
| 4 | Goat antirabbit IgG | HRP | ThermoScientific | 32460 | | AB_1185567 | 1:1000 |
| 5 | anti tubulin | | SigmaAldrich | T5168-100UL | B-5-1-2 | | 1:10000 |
